# Correlation in Domain Fluctuations Navigates Target Search of a Viral Peptide along RNA

**DOI:** 10.1101/2021.06.06.447299

**Authors:** Sangram Prusty, Raju Sarkar, Susmita Roy

## Abstract

Biological macromolecules often exhibit correlation in fluctuations involving distinct domains. This study decodes their functional implications in RNA-protein recognition and target-specific binding. The target search of a peptide along RNA in viral TAR-Tat complex is closely monitored using atomistic simulations, steered molecular dynamics simulations, free energy calculations, and a machine-learning-based clustering technique. An anti-correlated domain fluctuation is identified between the tetraloop and the bulge region in the apo form of TAR RNA that sets a hierarchy in the domain-specific fluctuations at each binding event and that directs succeeding binding footsteps. Thus, at each binding footstep, the dynamic partner selects an RNA location for binding where it senses higher fluctuation, which is conventionally reduced upon binding. This event stimulates an alternate domain-fluctuation which then dictates sequential binding footstep/s and thus, the search progresses. Our cross-correlation maps show that the fluctuations relay from one domain to another specific domain till the anti-correlation between that inter-domain fluctuations sustains. Artificial attenuation of that hierarchical domain fluctuation inhibits specific RNA binding. The binding is completed with the arrival of a few long-lived water molecules that mediate slightly distant RNA-protein sites and finally stabilizes the overall complex. The study underscores the functional importance of naturally designed fluctuating RNA motifs (bulge, tetraloop) and their interplay in dictating the directionality of the search in a highly dynamic environment.

## INTRODUCTION

From microbes to higher organisms, the symbiotic relationship between RNA and protein is the key to many cellular functions and regulations ^1–6^. Unfortunately, due to the unavailability of adequate structural information, compared to protein-protein or protein-DNA, studies exploring the RNA-protein recognition mechanism are limited. However, the intrinsic flexibility of both proteins and nucleic acids, perhaps, presenting one of the critical parameters helps to guide proteins to RNA and their binding in a sequence-specific manner^7–10^. To be more precise, the overall target recognition is governed by two fundamental processes: the efficient search and then the specific binding. Over the years, along with an interaction-level description (e.g., electrostatic, base-stacking, H-bonding), the profound role of conformational plasticity achieving strong specific binding upkeeps the famous “lock and key” binding metaphor^11–15^. While exploring the binding stability of a RNA-protein complex is high-rated as this complex eventually operates the required functions, the target search mechanism by which it is destined to form that complex is a priori important but less explored.

In the case of the DNA-protein complex, it has been well established that the search of protein for its binding site is guided by both one dimensional and three-dimensional search attributed to sliding and hopping^16–18^. The target search again involves a conformational proofreading mechanism^19^ where the conformation changes, unless optimal, would affect the fidelity of RNA binding by protein. This observation led to the extension of the induced-fit or conformational capture model initially developed by Koshland^11^ for enzyme-substrate binding to explain this phenomenon. The induced-fit model states that the first encounter of the protein with either RNA or DNA induces conformational changes in either or both of protein and RNA(/DNA) to form the biologically functional conformation of RNA-protein or DNA-protein complex^12,20^. This mechanism has been explored in RNA-protein complexes like HIV and BIV TAR-Tat ^21–24^ and Rev-RRE complexes ^25,26^. However, the induced fit model accounted for the specificity of the DNA-protein, or RNA-protein interaction was yet to illuminate a detailed mechanistic description of the target search and binding process. Meanwhile, a protein-centric mechanism known as the “fly-casting mechanism” was proposed by Wolynes and co-workers ^27^, which accounted for the shortcomings of the induced fit model. The “fly-casting mechanism” states that the unfolded protein binds the DNA weakly and non-specifically at a relatively large distance, which is subsequently followed by the folding of the protein as the protein comes closer and closer to its binding site^27,28^. This process is accompanied by a reduction in configurational entropy, an increase in negative enthalpy by forming stable interactions, and gain of entropy by solvents, thus providing an overall negative free energy change to the process^27^. A funnel-shaped energy landscape guides the search process of protein along the DNA similar to that of the free energy landscape of protein folding, having a greater global gradient towards the native structure as compared to that of local ruggedness ^27^. The role of conformational plasticity in RNA-protein recognition and the abundance of disordered regions in RNA-binding proteins (RBPs) have also been well established^14^. Moreover, it has also been found that these disordered regions in RBPs are highly conserved and make direct contact with the RNA for recognition^9^. Thus the flexibility of the binding partners is usually crucial for the RNA-protein recognition.

From the nucleic acids point of view, it has been seen that RNA has greater local flexibility than that of the DNA as it has a myriad of secondary structural elements such as bulges and loops. This flexibility of RNA has been reported to enhance the binding propensity of the protein to the RNA^10,29^. In some experiments, it has been observed that increasing the size of the bulge region of RNA leads to an increase in its binding affinity for the Tat protein as the flexibility of the RNA increases^30,31^. Jernigan and co-workers ^10^ have also shown using lattice model calculations that increasing the RNA bulge size leads to an increase in conformational entropy of the system and thus leads to an increase in affinity for Tat binding. This larger bulge entropies increase the probability of protein to bind to the RNA and thus aid in achieving the native bound conformation. Moreover, it has also been reported that asymmetric loops create strain in the RNA secondary structure ^32^ and thus leads to widening of major groove, facilitating the cognate protein to bind to the RNA^33,34^.

While all of these observations provide a general insight into the importance of conformational fluctuation of the binding partners, the more intriguing questions are: How the conformational changes/fluctuations can facilitate the molecular search? Whether these conformational changes hold secret signals that are sensed by the cognate partner in motion? Can this conformational fluctuation control the directionality of the search? This work investigates a detailed target search mechanism which not only involves site hopping and conformational proofreading it also shows how the conformational fluctuations direct the search path. Along with the search progress, the study also finds an intriguing role of some long-lived water mediators connecting the long-range specific RNA-protein sites. As a case study, we have investigated the RNA-protein complex formation process between TAR RNA and Tat peptide of Bovine Immunodeficiency Virus (BIV). TAR-Tat represents a structurally well-characterized RNA-protein model for understanding protein/peptide-mediated RNA target recognition mechanism. Tat is one of the crucial proteins encoded by immunodeficiency viruses, which binds to the trans-activation responsive(TAR) region of the RNA for transactivating the viral RNA, an essential step for viral replication ^35,36^. We have used atomistic simulation, steered MD, free energy calculations, and clustering techniques to explore the search pathway of the peptide by which it locates its thermodynamically stable RNA binding sites.

## METHODS

### System Preparation

The initial configuration of BIV TAR RNA bound to Tat peptide was obtained from the protein data bank(PDB ID: 1BIV) from a solution NMR structure reported by Patel and co-workers ^22^. The PDB was subsequently modified by removing the phosphate capping at the 5’-end of the TAR RNA to generate the initial structure of the BIV TAR-Tat complex for simulations. The initial structure of free BIV TAR RNA was obtained by removing the coordinates of the Tat peptide from the BIV TAR-Tat initial structure. All of the simulations were performed using the GROMACS 2018.3 package^37,38^. Topologies of all atoms were generated using the Amber 99 force field^39^ with parmbsc0^40^ and chiOL3^41^ modifications. The RNA-protein system and free RNA system were centered in a cubic box of length,12 nm, and were solvated with the TIP3P water model^42^. Subsequently, water molecules and sodium and chloride ions were added to both the systems to maintain the charge neutrality and mimic the system’s aqueous physiological environment. Details about the number of species of solvent and co-solvent are provided in Table S1.

### Simulation Protocol

After the complete system have been prepared, the potential energy of both the system was minimized using the steepest descent algorithm for the removal of steric clashes. Subsequently, for the complex system, both RNA and protein, and for the free system, only RNA was position restrained with force constant of 1000 kcal/mol/nm, and ions were frozen. After that, the system was equilibrated for 1 ns in an NVT ensemble to ensure proper hydration of the ions. The ions were then released at constant volume and were allowed to equilibrate for 3 ns. The RNA and protein were then slowly released by decreasing the force constant gradually from 1000 to 10 kcal/mol/nm^2^ at constant volume for 3 ns, followed by complete removal of position restraint on RNA and protein at constant pressure for 500ps. Hereafter, unrestrained explicit solvent simulations were carried out for 200 ns in the NVT ensemble. Throughout all our simulations, we used a leapfrog integrator with a time step of 2 fs and maintained an average temperature of 300 K using a Nose-Hoover thermostat^43,44^ with a relaxation time of 0.5 ps. For all the NPT equilibration along with the above-mentioned temperature conditions, we used Parrinello-Rahman barostat^45^ to maintain an average pressure of 1 bar with a coupling time constant of 0.5 ps. Periodic boundary conditions were used in all the simulations in all directions. The neighbor list was updated after every 10 steps using a grid system with a short-range neighbor list cutoff of 1 nm. Particle Mesh Ewald(PME)^46^ was used to calculate the electrostatic interactions with a Fourier spacing of 0.12 nm and an interpolation order of 4. All bonds were constrained using the LINCS algorithm^47^.

### Constant Velocity Steered Molecular Dynamics Simulations

The unbinding event associated with the dissociation of a protein from the RNA requires energy greater than that of the thermal energy at room temperature. Therefore we cannot access all the states associated with the unbinding event using standard molecular dynamics simulations. So, we used a constant velocity steered molecular dynamics(cv-SMD) approach in which a time-dependent external force is applied to the protein along the reaction coordinate to facilitate the unbinding. Specifically, this unbinding event is captured by adding an extra harmonic bias potential along the reaction coordinate to the hamiltonian. Subsequently, the protein is pulled away from the RNA with a constant velocity^48,49^.

The cv-SMD simulations were also performed using GROMACS 2018.3^37^ package. The distance between the center of mass of TAR RNA and the center of mass of Tat peptide was chosen as the reaction coordinate for pulling the Tat peptide out of its binding site in the TAR RNA. A force constant of 500 kJ/mol/nm^2^ and a pulling speed of 0.001 nm/ps was used for the simulation.

### Free Energy Calculation: Umbrella Sampling

The free energy difference between two states A and B can be evaluated using the following expression:

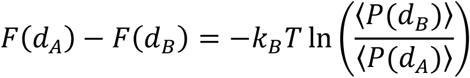

where 〈*P*(*d_A_*)〉 is the probability of obtaining the system in state A at reaction coordinate *d_A_*, and 〈*P*(*d_B_*)〉 is the probability of obtaining the system in state B at reaction coordinate *d_B_*. But this expression cannot be used for calculating the free energy if a high energy barrier separates the two states due to poor sampling of one of the states. So, to tackle this problem, umbrella sampling^50^ is widely used.

For the calculation of the free energy profile also, the distance between the center of mass of the RNA and the protein is taken as our reaction coordinate. And we used the trajectories generated by cv-SMD simulations to generate initial configurations for each umbrella sampling windows. The starting configuration for 29 umbrella windows was chosen based on the center of the mass distance of the RNA and the protein from this trajectory. We used an asymmetric way to choose our sampling windows to get overlapping umbrella windows. The overlap of the windows is shown in Figure S1. Each of the extracted configurations for the umbrella sampling windows was simulated for 10 ns in the NVT ensemble at 300K with a force constant of 500 kJ/mol/nm^2^ and 0 nm/ps pull speed. The average temperature was maintained using a Nose-Hoover thermostat^43,44^. After this, we used the weighted histogram analysis method(WHAM)^51^ to combine all the individual potential of mean force(PMF) to generate the unbiased, free energy profile.

### Simulation for the association of TAR-Tat complex

This work aims to investigate the binding mechanism of the TAR RNA to that of the Tat peptide. Thus to look at the binding mechanism, different configurations were extracted from different time frames from the SMD trajectories. Multiple unbiased molecular dynamics simulations were performed using the same parameters as used in unrestrained explicit solvent simulations, as mentioned previously. A set of unrestrained explicit solvent simulations were necessary for designating landmark states along the binding pathway.

### Analysis of Simulation Trajectories

To have a more in-depth insight into the binding pathway of Tat peptide with the TAR RNA, we have calculated the root mean square deviation(RMSD), root mean square fluctuation(RMSF), the evolution of native contacts and native contact maps, dynamic cross-correlation maps, residence time distribution. We have also done a principal component analysis(PCA) to understand the anti-correlated motion of the bulge and the tetraloop. A detailed description of these analyses is provided in the *Supporting Methods*.

## RESULTS AND DISCUSSION

### Free energy profile and preferential binding pathway of TAR-Tat interaction

Tat is well recognized for its Arginine-Rich Motifs (ARMs) ^22,35^. These ARMs are quite potent. Being doubly positive charged, they efficiently interact with the negatively charged phosphate backbone of RNA and also forms non-canonical stacking interactions with RNA bases, which helps Tat to specifically bind TAR with a high binding affinity^52–54^. Understanding the interactive forces that drive Tat to attain the high binding affinity and specificities over other non-specific interaction-based noises requires the integrated knowledge of binding and conformational breathing of both the binding partners.

The binding is monitored from the free energy profile as a function of the distance between the center of mass of the TAR RNA and the Tat peptide describing BIV TAR-Tat complex formation from the apo-form of BIV TAR RNA and Tat peptide (**Figure 1C**). The profile shows that the Tat peptide binding to the TAR RNA is entirely a downhill process only with a few humps where the slope of the barrier is observed to change noticeably. Based on these positions where there is partial binding and change in the barrier slope, four different binding states, E, D, C and B are defined to characterize the binding progress from their free form to the bound form, respectively as shown in Figure S2. The global minimum is defined as state A, (representative structures of all the binding states are shown in Figure S3-S8).

**Figure 1.**
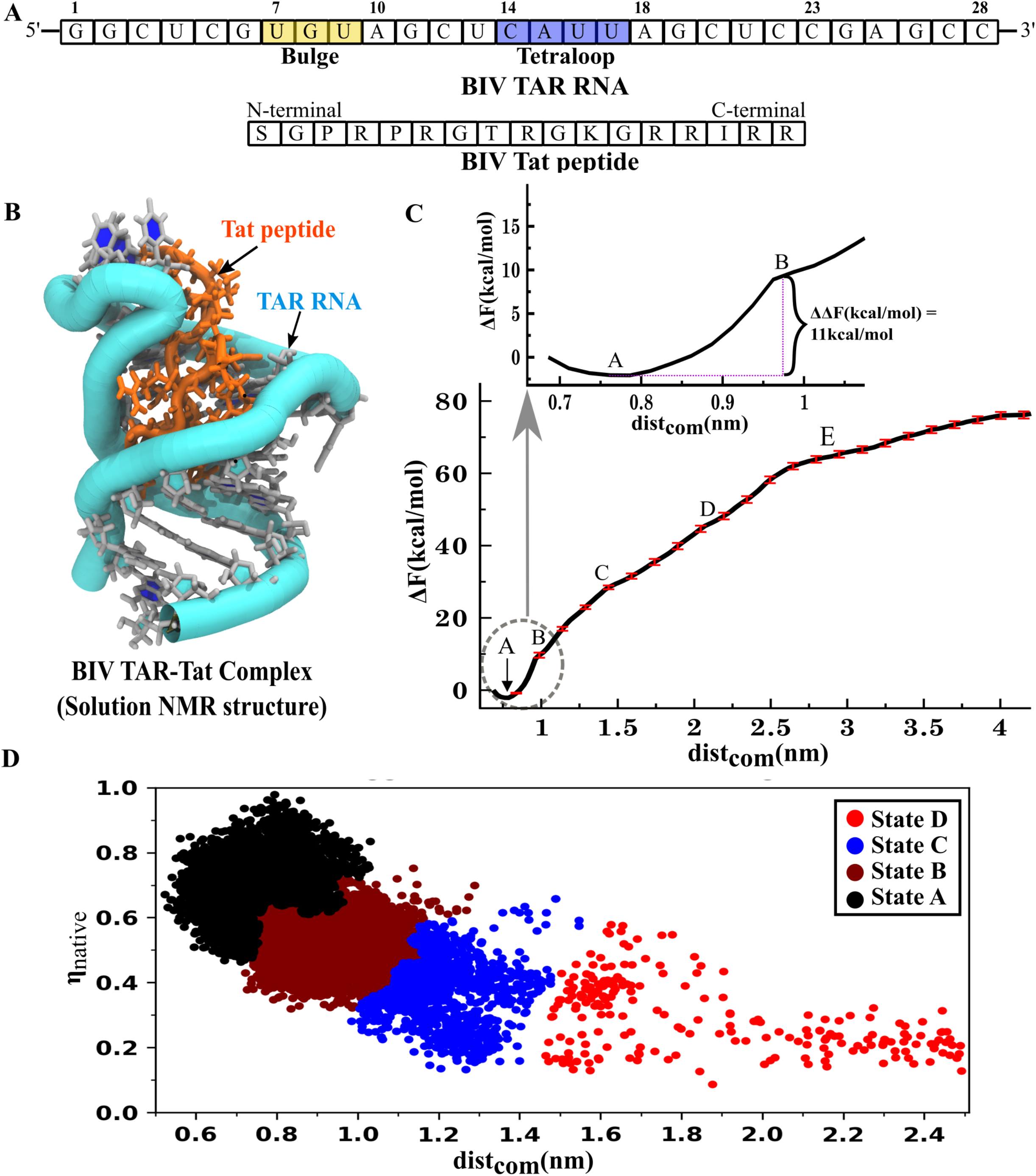
Binding pathway of the formation of BIV TAR-Tat complex. (A) Primary structure of BIV TAR RNA and BIV Tat peptide. The bulge and the tetraloop region of the BIV TAR RNA are highlighted in yellow and blue color, respectively. (B) All-atomistic conformation of BIV TAR-Tat complex as taken from a solution NMR structure (PDB id: 1BIV). (C) Average free energy profile associated with the binding of BIV TAR RNA and BIV Tat peptide for the formation of BIV TAR-Tat complex. The states associated with the binding progress are labeled as E, D, C, B, and A The position marked as A corresponds to the native conformation of the BIV TAR-Tat complex. The free energy loss(11kcal/mol) involved in the final, binding phase is shown in the inset figure. (D) Scatter plot depicting the dominant binding pathway and the presence of four clusters (defined based on Ward’s agglomerative clustering method) in the binding pathway of TAR-Tat complex based on fraction of native contact(η_native_) and dist_com_.

The change in free energy involved in specific binding (**inset of Figure 1C**) is calculated to be ~11kcal/mol. A similar result has also been obtained in the isothermal titration calorimetry experiment by Goel et al.^55^, where they obtained a free energy difference between the bound and unbound state of TAR RNA to be ~9.5kcal/mol. The small discrepancy between experimental and computational calculations might arise due to a known sampling issue of the unbound form. This quantification error range doesn’t harm the purpose of exploring the search pathway. However, it is also observed that as the binding of the protein on the RNA progresses, there is a gradual increase in the slope associated with the successive binding steps (Table S2 and Figure S2). These slopes characterize the stiffness of the barrier separating subsequent energy states. Thus, the increase in the slope with binding progress suggests a possible cooperative binding mechanism where each binding step facilitates the next binding move, synonymous with the cooperativity present in the protein folding pathways ^56,57^.

Each state in this binding pathway is stabilized by several non-covalent interactions such as hydrogen bonding^58,59^, electrostatic^54,60,61^ and base stacking interactions^59–62^ occurring between the nucleotide of the TAR RNA and the amino acid residues of the Tat peptide^52–54^. These electrostatic and base stacking interactions are majorly mediated by the guanidium group of several arginine residues of the Tat peptide^26,30,63,64^. A detailed molecular picture of the significant non-covalent interactions involved in each landmark state (E to A) along with a newly found ARM is discussed in *Supporting Results and* Figure S4-S8.

While the changes in the slope in the 1D free energy profile efficiently identifies the landmark states, the information related to their conformational dynamics during the binding progress is too limited as the sampling along a single reaction-coordinate covers a restricted phase-space. However, the binding and binding induced conformational changes occur in higher dimensional phase space. To investigate such induced conformational changes along the preferential binding pathway, we have sampled each state (D to A, essentially describing peptide’s sojourn to its target binding location) of the RNA-protein complex extensively and repetitively using unrestricted atomistic MD simulations (see ***Methods***). After the preliminary designation of different landmark states and identifying their initial configurations, in the next assignment, as the binding pathway is downhill it was straightforward but instrumental for us to generate multiple sets of long unbiased association trajectories. From all these association trajectories, we have identified our early designated states and their survival time. Within this survival time, an ensemble of close configurations was generated for each designated state. This ensemble of configurations is projected over an order parameter plane of the fraction of native contacts (*η_native_*) and the center of the mass distance between the TAR RNA and the Tat peptide (dist_com_) and Agglomerative Ward’s clustering is then employed to obtain the cluster of population states connecting the entire binding pathway as shown in **Figure 1D**. The 4 clusters formed have a similar center of the mass distance between the TAR RNA and the Tat peptide (dist_com_) as obtained for the state D, C, B, and A from the 1D free energy profile. However, in state E, both RNA and protein are in their apostate.

We have also used a steered molecular dynamics (SMD) simulation approach to sample the phase space capturing the unbinding event by applying a bias potential and pulling the Tat peptide away from the TAR RNA (See the ***Methods*** section for detailed steered MD simulation method used). From the ensemble of ten independent SMD simulations, we observe the population convergence of association and dissociation pathways from multiple trajectories scanning the overall binding phase space (Figure S9). In a few trajectories, a backtracking phenomenon has been observed just before the final, binding step during the transition from step B to native state A in which there is a sudden loss of native contacts, and then the native contacts are again gained as the native state A is approached (Figure S10). A similar backtracking event has also been documented in protein folding pathways ^65^ and multidomain protein association ^66^.

### A relay in fluctuation induced binding and binding induced fluctuation dictates the binding pathway

The conformational fluctuation-induced binding progress and binding-induced conformational fluctuation are monitored from state-D to state-A, sequentially. The evolution of conformational fluctuation over the binding in progress are quantified and connected by calculating the root mean square deviation (RMSD) distribution of bulge and tetraloop regions of BIV TAR RNA and RNA-protein intermolecular contact map. The connection is illustrated in **Figure 2**. The epicenter for the tetraloop and bulge fluctuation is present in the tetraloop and the bulge respectively however these fluctuations exert local effect on their neighboring nucleotides also. Thus the RMSD is calculated on TAR RNA residue 8-13 for bulge fluctuation and for tetraloop fluctuation residue 15-20 are taken into account. These two regions are named as the “near tetraloop region” and “near bulge region” to account for both the epicenter and local spread of fluctuation. The native contact map is derived at an atomistic level to augment the count. For reading convenience, the atomistic and nucleotide-residue level sequence is shown in **Figure 2A** (TAR) and **Figure 2B** (Tat). The native contact probability greater than 60% is considered to plot the native contact map for each binding state. The pattern and magnitude of fluctuation at the RNA residual level are observed to influence the peptide’s sequential binding footstep.

**Figure 2.**
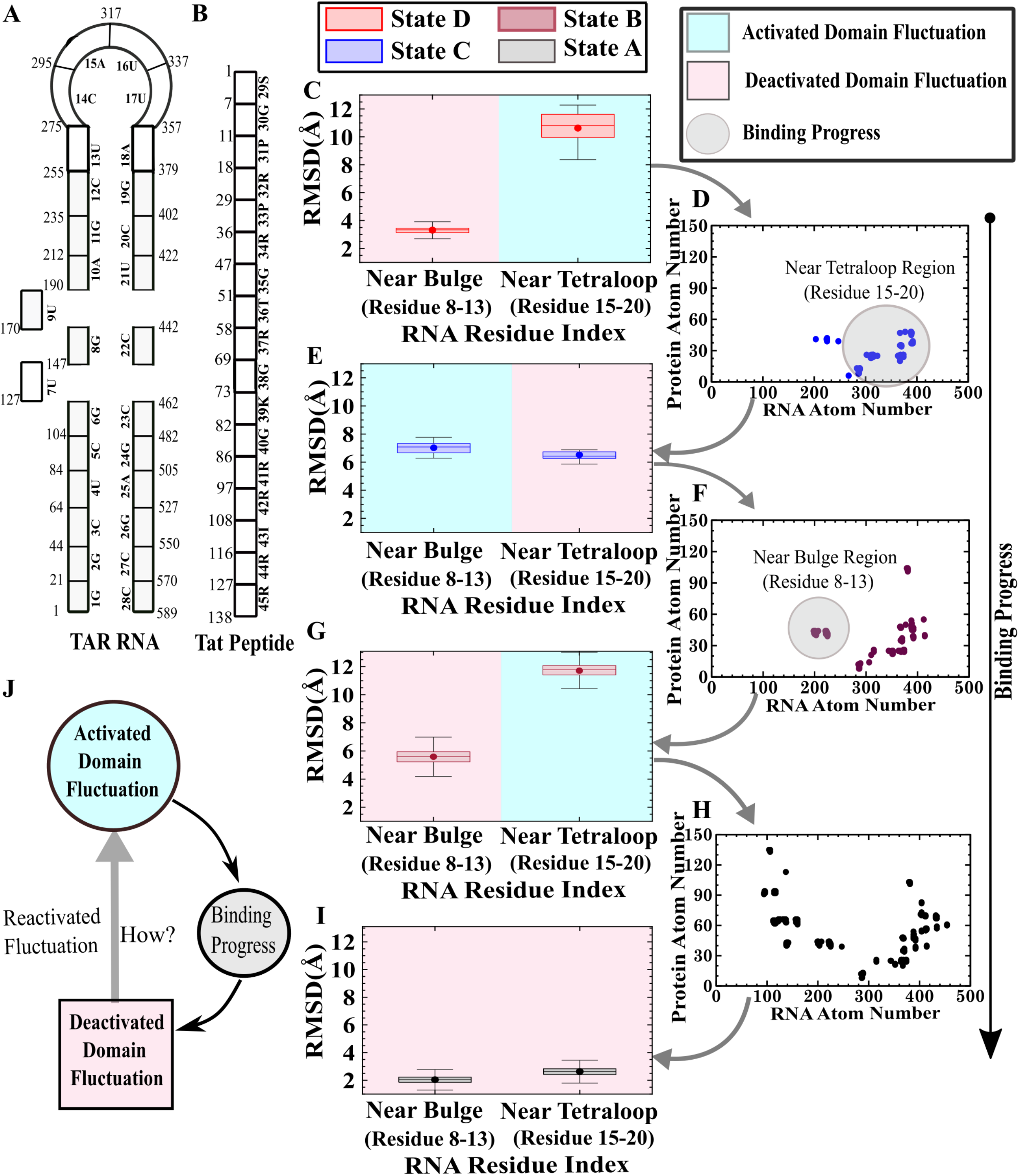
The cycle of activation of domain fluctuation and its binding induced dissipation. (A) Secondary structure of BIV TAR RNA along with the atomic index (shown in non-bold numerals) corresponding to the residue index(bold numerals written along with the nucleotide symbol). (B) The sequence of BIV Tat peptide along with the atomic index (shown in non-bold numerals) corresponding to the residue index(bold numerals written along with the amino acid one letter symbol). (C) Boxplot of Root means square deviation (RMSD) for near bulge and near tetraloop region of state D showing the activated(higher) fluctuation of near teraloop region. (D) RNA-protein atomistic contact map for binding state C depicting native contact formation around tetraloop region of BIV TAR RNA. (E) Boxplot of Root means square deviation (RMSD) for near bulge and near tetraloop region of step C showing the activated and deactivated fluctuations of near bulge and the near tetraloop region respectively. (F) RNA-protein atomistic contact map for binding state B depicting native contact formation around the bulge region of BIV TAR RNA. (G) Boxplot of Root means square deviation (RMSD) for near bulge and near tetraloop region of step B showing the reactivated and deactivated fluctuations of near tetraloop and the near bulge region respectively. (H) RNA-protein atomistic contact map for native state A depicting all the native contact formation. (I) Boxplot of Root means square deviation (RMSD) for near bulge and near tetraloop domain of step A showing the complete deactivation of fluctuation around both the tetraloop and the bulge due to complete native contact formation. (J) Flowchart depicting the cycle of activation of domain fluctuation followed by binding induced deactivation and domain fluctuation transfer. Note that the native contact probability greater than 60% is considered to plot the native contact map for each binding state. In the boxplot, the box corresponds to the 25-75 percentile of the distribution and the dot and line inside the box represents the mean and median of the distribution.

The binding footsteps progress with the following events:

i. In the apo form of RNA, the near tetraloop region (residue 15-20) shows higher fluctuation as in state-D than that of state-C (**Figure 2C**).
ii. At an initial approach, the Tat peptide makes a few binding contacts around the near tetraloop region and binds partially (**Figure 2D**).
iii. This partial binding causes a significant reduction/deactivation in near tetraloop region fluctuation (**Figure 2E**). The detailed information of the conformational changes and intermolecular interaction is illustrated in Figure S6, where the leading role of ARMs in this search process is well captured. The ARMs associated binding influences such a conformational change that it leads to the activation of fluctuation in the near bulge region (residue 8-13) in state-C, as shown in **Figure 2E**.
iv. Once again, the peptide senses the higher fluctuation reflected in the RMSD distribution promoting the binding event occurring at the near bulge region with a number of newer native contacts in state-B (**Figure 2F**).
v. The pattern relays with the binding induced deactivation of individual domain fluctuation and the activation/reactivation of an alternate domain fluctuation. Thus, in state-B, near bulge binding leads to the deactivation of near bulge fluctuation and the reactivation of near tetraloop fluctuation (**Figure 2G**).
vi. At this point, the intermolecular RNA-protein contact map shows the formation of a few other native contacts in their overall interfacial regime (**Figure 2H**), and the complex acquires nearly 95% of intermolecular native contact generation.
vii. This late-stage formation of native contacts doesn’t cause any more fluctuation change to the alternate domain. It instead reduces the magnitude of fluctuation of the overall complex (**Figure 2I**).

Correlating all the above observations in **Figure 2J**, we illustrate the cycle of conformational fluctuation induced binding and binding induced fluctuation deactivation and transduction. This cycle summarizes that at each binding footstep, the peptide senses the hierarchy in fluctuation generated by a set of nucleotides of the RNA. For binding, it selects that specific domain/nearby region preferentially, which has the activated/higher fluctuation. We observe suppression of fluctuation/dynamics which is quite expected upon binding. However, which is hard to intuit from any past binding event is the understanding of binding induced conformational changes and, as a result of these changes, which domain fluctuation will be enhanced next from its previous value to navigate the binding. Thus, how reactivation of an alternate domain fluctuation stimulated is a legit question here.

Before we explore the origin of reactivated fluctuation, to further substantiate on hierarchical fluctuation mechanism in terms of fluctuation activation and deactivation in the TAR-Tat binding pathway, we monitor the difference between the root mean square fluctuations (RMSF) of adjacent binding state of each nucleotide of the BIV TAR RNA. The RMSF is calculated on the phosphate atom of each nucleotide for clear analysis. Analysing the RMSF difference between state D and state C (**Figure 3A**) it was clear that at state D, the near tetraloop region has an activated fluctuation which gets deactivated in state C 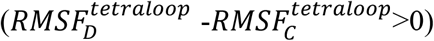 and activation of near bulge region fluctuation occurs in state C 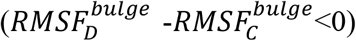. Subsequently in state B, there is deactivation of near bulge fluctuation 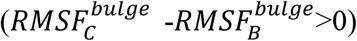 and activation of near tetraloop fluctuation 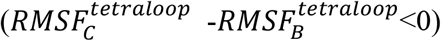 as shown in **Figure 3B**. Finally when the TAR-Tat complex acquires its native state conformation (state A) there is only decrease in fluctuation between domains 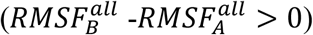 as shown in **Figure 3C**. The raw RMSF plots for each of the binding states is provided in Figure S11. Moreover, the role of fluctuation from peptide’s perspective in the binding pathway has been discussed in Supporting Results and Figure S12.

**Figure 3.**
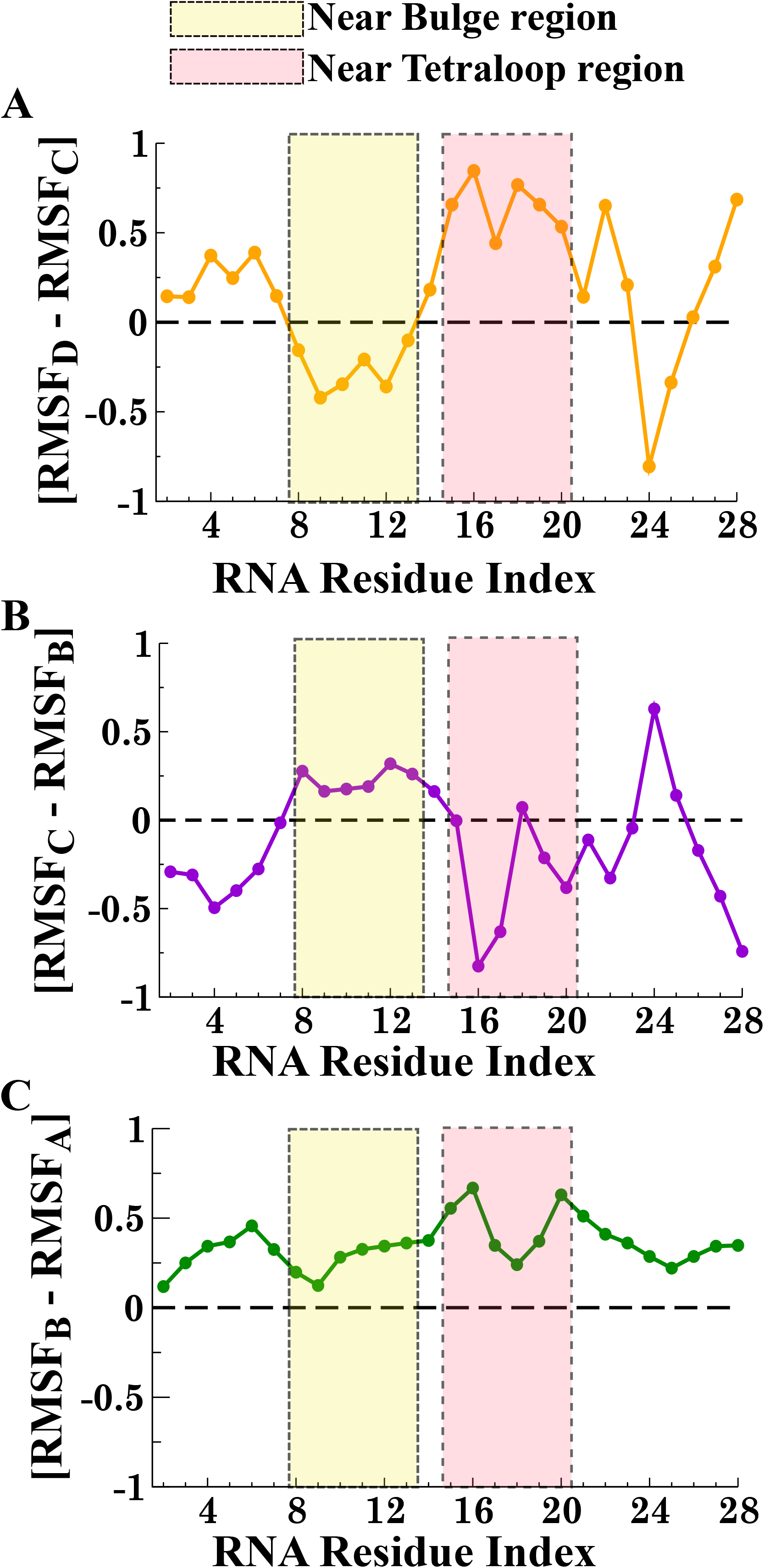
Hierarchical fluctuation between the near tetraloop and the near bulge region of TAR RNA is followed by measuring the difference between the RMSF (Root Mean Square Fluctuation) of successive binding states. (A) RMSF difference between state D and state C of binding showing an activated fluctuation in near tetraloop region which gets deactivated in state C 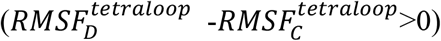 and subsequently fluctuation activation occurs at near bulge region in state C 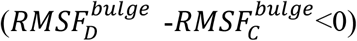. (B) RMSF difference between state C and state B of binding showing deactivation of bulge fluctuation 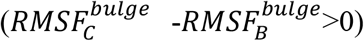 and activation of tetraloop fluctuation 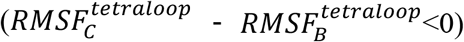 in state B. (C) RMSF difference between state B and state A showing only decrease in fluctuation from state B to state A 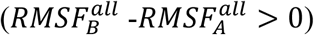.

### Anti-correlation in alternate domain fluctuations regulates hierarchical fluctuation

We urge to understand an interesting observation that we followed in **Figure 2** and **Figure 3** which is the following: We realize that suppression of fluctuation of a certain domain is quite expected upon its association with its binding partner. However, while fluctuation of one domain/region becomes suppressed/deactivated, we interestingly observe an alternate specific domain to enhance/reactivate its fluctuation than the fluctuation in its previous state (**Figure 3**). Does that alternate domain complement the fluctuation-deprived-domain to maintain the hierarchical fluctuation and thus to direct the pathway? If it does so, we anticipate that these two complementary domains may show certain correlation in terms of their fluctuation. To find the connection between near bulge and near tetraloop region the correlation between domain-domain fluctuations is quantified at each binding step using dynamic crosscorrelation analysis. Details about the calculation of dynamic cross-correlation have been provided in the *Supporting Methods* section. The cross-correlation map for apo TAR RNA and four TAR-Tat binding states: D, C, B, and native state A is presented in **Figure 4**. The cross-correlation map for state D shows that the tetraloop of the TAR RNA is highly anticorrelated to that of its bulge region. The same anti-correlation was also observed in state C; however, the correlation coefficient is slightly lowered than that of state D. Subsequently, with the progress in binding, the anti-correlation between the tetraloop and bulge region is gradually faded. Moreover, in the apo state, also there is the presence of a feeble anticorrelation between the bulge and the tetraloop region which then gets amplified in state D on initial approach of Tat peptide to the TAR RNA. This anticorrelation present between the bulge and tetraloop has been observed to initiate the hierarchical search by keeping the fluctuation of tetraloop higher and fluctuation of bulge lower in the initial association event which leads to initial binding at the tetraloop region and subsequently upon binding-induced suppression of tetraloop fluctuation, bulge fluctuation increases due to this prevalent anticorrelation and protein then binds at the bulge region. Furthermore the other anticorrelation present in the dynamic cross correlation map corresponds to highly flexible terminal RNA residues which have the inherent characteristic of flexibility.

**Figure 4.**
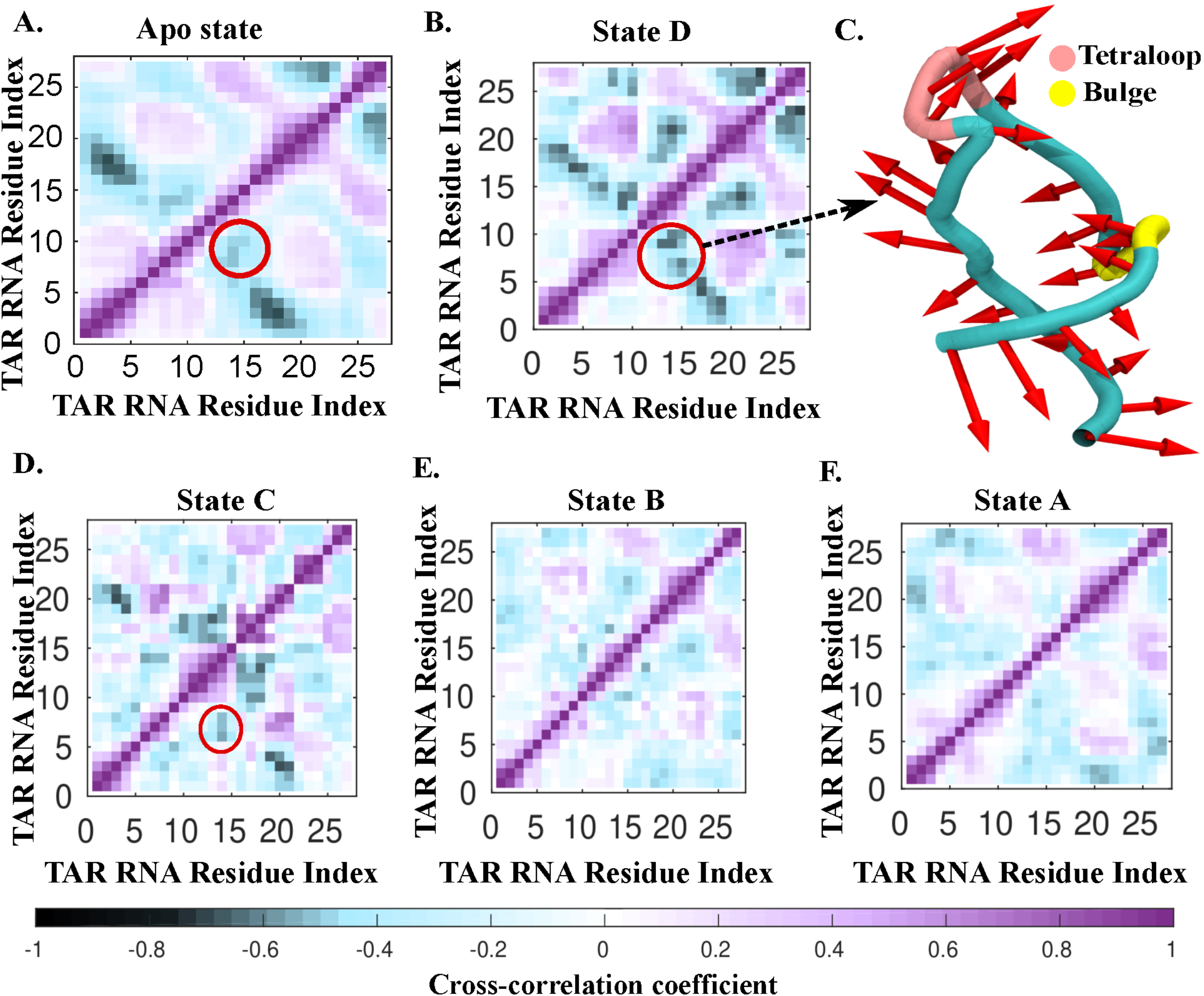
Non-linear progression of anti-correlation in the fluctuation of dynamic tetraloop and bulge region with the binding-progress. (A) Cross-correlation between the fluctuation of the phosphate atom of each of the residues of the TAR RNA calculated on apo state. A feeble anticorrelation between tetraloop and bulge is present in this state shown with red circle. (B) Cross-correlation between the fluctuation of the phosphate atom of each of the residues of the TAR RNA calculated on state D showing amplified anticorrelation between tetraloop and bulge depicted with a red circle. (C) Second PC mode of state D conformation of the TAR RNA calculated on phosphate atom of each nucleotide, capturing the tertraloop-bulge anti-correlated motion (D) Fading of anticorrelation of bulge and tetraloop in state C(shown in red circle) (E) Cross correlation between the TAR RNA residue in state B showing nearly further fading of bulge-tetraloop anticorrelation (F) Cross correlation between the TAR RNA residue in state A showing low anticorrelation between the bulge and tetraloop.

To further substantiate the anti-correlated motion of the tetraloop and the bulge, we have used elastic network modeling to generate the slow modes of the correlated motion associated with each of the binding states ^67^. The principal component analysis beautifully captures the directionality of this anti-correlated domain motion highlighted in state-D (**Figure 4C**). Directionality of anti-correlated motion for the other states (C to A) is shown in Figure S13.

### Artificial attenuation of tetraloop fluctuations inhibits binding

We have observed that the correlation that exists between tetraloop (TL) and bulge region sets a hierarchy in the corresponding region-specific fluctuation at each binding step and that may bias the binding pathway. However, correlation between alternate domains may not imply causality. One need to check that if fluctuation of one of those correlated domains is artificially attenuated, would binding be inhibited? To investigate such possibility we employed unbiased molecular dynamics simulation method to simulate two situations/conditions starting from same configuration extracted from SMD trajectory. They are the following: (i) Restrained TL simulation: here position of referred tetraloop domain is weakly restrained and (ii) Unrestrained TL simulation: here no restraining force is applied on the position of TL domain. For the above two conditions, the time evolution of the fraction of native contact was calculated from their respective trajectories. On attenuation of TL fluctuation, we find that the binding is certainly inhibited there while TL was the initial binding target (**Figure 5**). We have characterized both the prevalence of fraction of native contact and also all-level contacts (accounting for native and non-native contact) in the TL (Figure S14) which substantially decreased from its earlier situation when it was guided by its natural fluctuation. Instead, as the relative fluctuation of the bulge is now higher than that of the tetraloop, the protein targets bulge as its initial binding target as shown in Figure S14.

**Figure 5.**
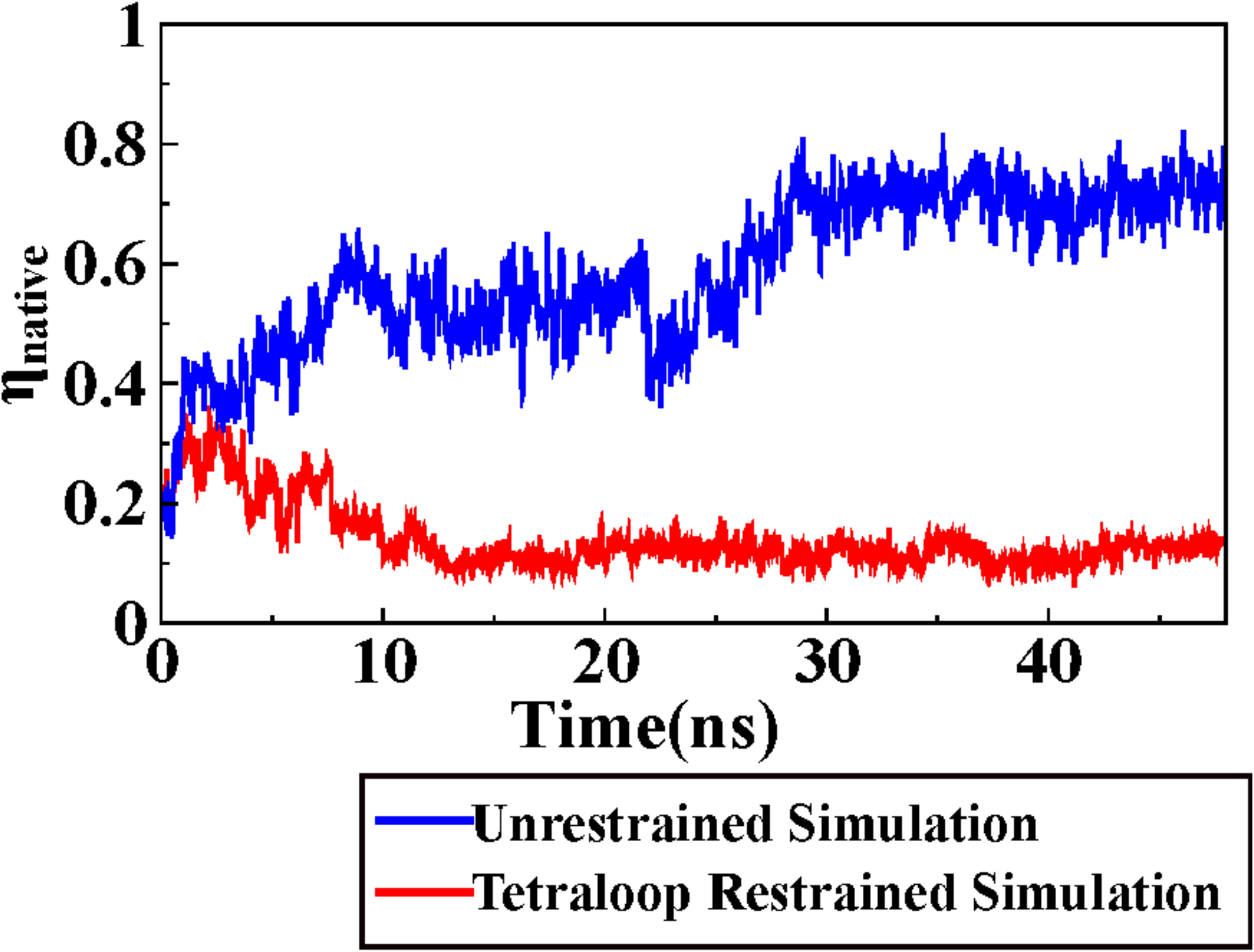
Binding of Tat peptide to TAR RNA facilitated by the specific nucleotide fluctuations investigated from time evolution of fraction of native contact. Time evolution of fraction of native contact has been calculated on two simulation trajectories one with position restraints on the near tetraloop region and another without any position restraints.

Summarizing all the above observations, we can conclude that at each binding step there is an activated domain fluctuation which is sensed by the peptide and it binds to that domain. As the binding occurs that domain gets deactivated and another specific domain’s fluctuation arises and this cycle continues till the protein finds a compatible and stable binding site. This finding is indeed consistent with early experimental observations by Weeks *et al*. ^31^ and Chen *et al.^30^* and theoretical assessment by Lustig *et al*. ^10^, where they found the larger bulge entropy to promote competent binding in the case of TAR-Tat complex. However, the overall recognition and binding is not a single step process involving only one domain. The dynamic partner (here peptide) moves between sites and scan for favorable interactions spanning the entire RNA to minimize the overall binding free energy. During this scanning process, whether fluctuation-deactivation of a particular domain correlates with another domain’s fluctuation-activation is an intriguing question here as this stands an opportunity to understand fluctuation induced signal transduction in molecular recognition.

### Water: The molecular glue finally firms the formation of TAR-Tat complex

The fluctuation pattern of BIV TAR RNA nucleotides for both state B and native state A are very similar, and only the magnitude of fluctuation of native state A is lower. The question arises, what drives this stabilization in the native state, A? We have investigated the dynamic coupling between water molecules and RNA sites during the binding progress by calculating the residence time distribution of water molecules in binding state (B, A and apo-form). Due to the transient nature of state D and C they are not considered for the residence time calculation.

Early time-resolved fluorescence measurement study performed by Maiti and coworkers on the same BIV TAR-TAT system showed that a magnitude of 1.8 ns in the solvent correlation time arises due to the hydration layer composed of water molecules hydrogen-bonded to the RNA bases or due to the diffusion of interfacial water molecules between the bound hydration layer and bulk water^55^. Based on this experimental finding, we have considered water molecules staying beyond 2 ns can be classified as long-lived bound water molecules. Thus, calculating residence time distribution we have classified bound-water molecules into three categories: water molecules having residence time (i) 2-4ns (ii) 4-6ns (iii)>6ns. ^68–70^ State B and A are found to be associated with a few quasi-bound water molecules that are motionally restricted with a long residence time of 2-6ns or more (**Figure 6**). The residence time distribution was calculated by taking a cutoff of 4.5Å, which corresponds to the water molecule’s second hydration shell around the BIV TAR RNA from the radial distribution of the water molecule around the TAR RNA (Figure S15). Details about the calculation of residence time distribution have been provided in the *Supporting Methods* section. The residence time distribution profile for the apo state of TAR RNA shows a peak of around 0.25 ns as shown in Figure S16. As the apo form associates with the peptide, the peak shifts towards higher residence time (Figure S16). This implies that TAR-Tat binding leads to a tighter solvent cage around the TAR RNA, due to which some water molecules have more restricted motions. Similar result have also been found by Maiti and coworkers^55^ experimentally where they obtained a longer timescale solvent relaxation of 5.3ns for BIV TAR-Tat complex.

**Figure 6.**
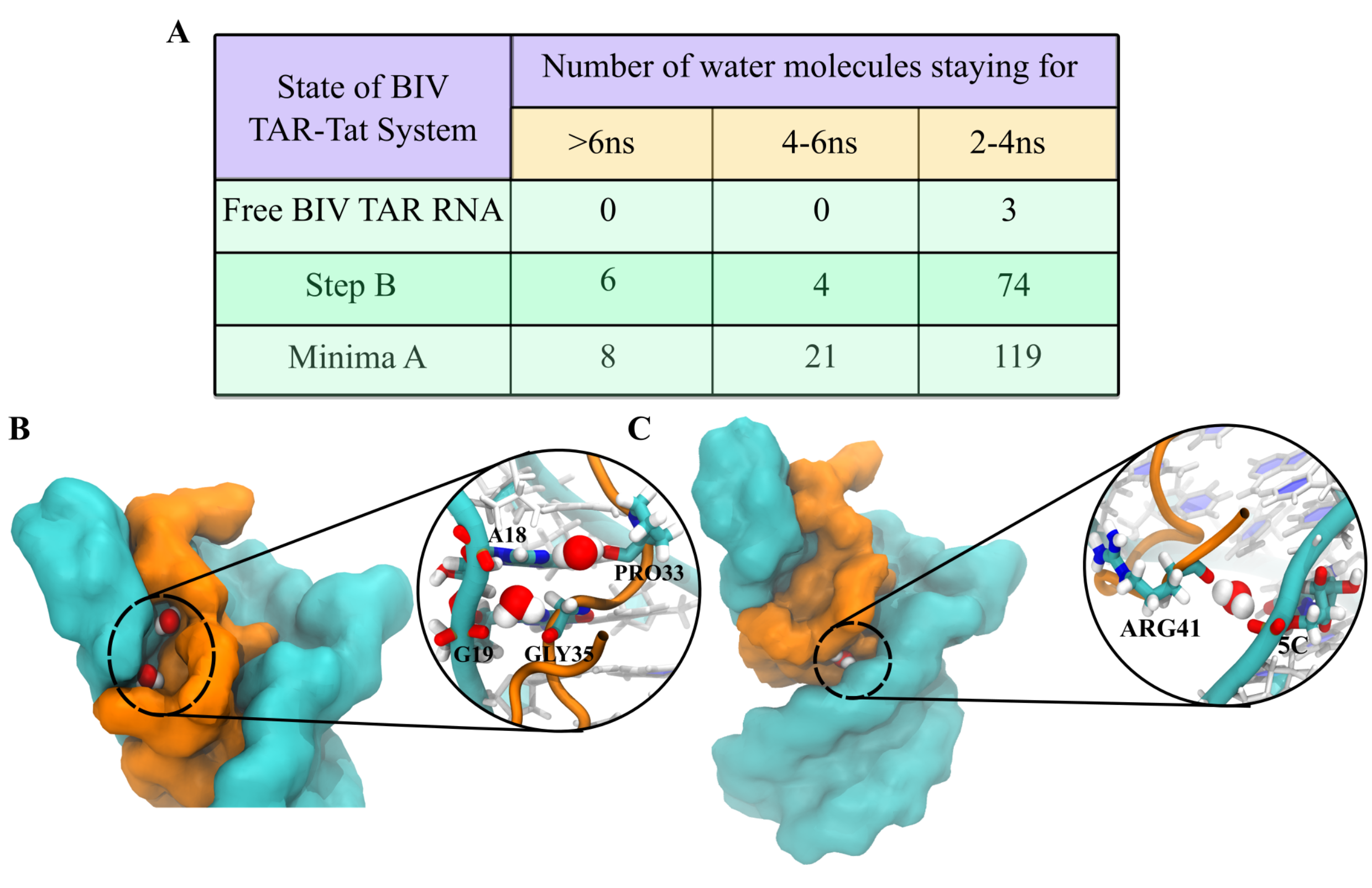
Water, the molecular glue for the formation of the native TAR-Tat complex. (A) The table categorizes the longer staying water molecules into three categories: water molecules staying for (i) >6ns (ii) 4-6ns (iii) 2-4ns. Based on these three categories, the number of water molecules found in each category at three different states of the BIV TAR-Tat system (free TAR RNA, Step B, and native state A) is recorded in the table. It was found that as we move from free BIV TAR RNA to step B and subsequently to native state A, the number of water molecules having high residing time increases. (B) Two water molecules having high residence time acting as a bridge between the TAR RNA and Tat peptide. One of the water molecules is mediating the interaction between the guanine and glycine residue, and another water molecule is mediating the interaction between the adenine and proline residue. (C) Arginine and cytosine backbone are interacting with the help of a highly residing water molecule.

The long-lived water molecules are grouped together as they belong to a specific residence time zone (**Figure 6A**), both for state B and state A. It is evident from this table that the water molecules indeed are acting as a mediator for stabilizing the TAR-Tat complex in the native state. An increase in the number of longer staying water molecules renders a decrease in the fluctuation of native state A compared to state B. Subsequently, the frequently visited most probable water-wetted sites are also detected. Interestingly, in state A, the longer staying water molecules are mostly found to mediate the interactions between those sites (e.g., G19-GLY35, A18-PRO33) where direct electrostatic interactions are not feasible. A snapshot of these water-mediated interactions is presented in **Figure 6B** for State A. For state B, it is shown in Figure S17.

## CONCLUSIONS

Our investigation was stimulated by a review article by Draper summarizing the consensus on RNA-protein specific recognition ^72^. The fact is touched upon that the structural details of RNA-protein complexes are being resolved by NMR and X-ray crystallography, but thorough thermodynamic analyses of recognition mechanisms have yet to be performed. Toward this prevailing direction, ours is certainly not the first investigation ^10,52,73,74^, but this effort brings upon a thorough thermodynamic analysis comprehending each step of the recognition and binding and their processing. The quest is to explore and understand the physical principle behind specific target recognition phenomenon if exists. The investigation is mainly driven by the long-standing questions in this field: How does the protein navigate its specific target location? What drives the specific binding of a protein to that of the RNA? In colloquial words, this first question still plea for understanding the recognition mechanism at an atomistic level, and the second one can be explained by the thermodynamics of binding. Nonetheless, both recognition and binding involve many degrees of structural changes. What is essential is to understand whether such structural changes imply a code/principle illuminating an efficient search mechanism. However, in this study, the efficiency of the search mechanism yet not implies a faster target search mechanism; it only indicates a possible guiding principle facilitating the search.

During the investigation of the specific RNA-protein recognition mechanism, we find the following connected aspects: (i) Higher conformational freedom and fluctuation provide a higher competence for binding (**Figure 2**). This phenomenon has also been observed in various experiments ^30,31^ and fairly explained in early theories^10^. Attenuating such specific conformational fluctuation may inhibit binding and alter its pathway (**Figure 5**). (ii) While binding deactivates the fluctuation of a specific domain, in turn, an alternate domain fluctuation turns activated (**Figure 2 and 3**). (iii) The activation of alternate domain fluctuation appears as a response to a prevailing anti-correlation between specific domain-domain fluctuations (**Figure 4**). This anti-correlation was already present in the apo-form of RNA but the response becomes amplified when protein approaches RNA vicinity (**Figure 4**). This study bolsters the fact that how domain-domain fluctuation correlation/anti-correlation induced hierarchical fluctuation pattern may be exploited by the system purposefully for signal transduction and binding site selection. The anti-correlated fluctuation-induced signal transduction eventually guides the dynamic partner, preferably towards the direction of minimum energy, and thus, the target search may become rather efficient. It is noteworthy that the proceedings of recognition are highly dependent on its hydration and ion environment. Water and ions stay highly dynamic during the process of recognition until a few long-lived water molecules mediate the final locking of the RNA-protein association in a form of the thermodynamically stabilized complex (**Figure 6**).

In conclusion, plenty of experimental and computational studies observe such dynamic, allosteric coupling between domains in a single biomolecule. With the growing pieces of evidence of highly dynamic biomolecules like RNA, disordered proteins, and the myriad of their diverse structural motifs, though highly challenging, it’s time to identify whether the coupling between these structural motifs indicates a meaningful functional signal veiled within complexity. In the present case of the TAR-Tat complex, it is evident that the dynamic coupling between two irregular and flexible RNA motifs, bulge, and tetraloop, performs the signal transduction for binding. The follow-up research will focus on the natural existence of these irregular motifs, whether deliberately designed to promote such recognition signals, in general. The investigation is underway in this direction.

## ASSOCIATED CONTENT

### Supporting Information

Supporting information is available free of charge via the Internet at http://pubs.acs.org.

➢ Supporting Methods: Tetraloop Restrained Simulations, Bulk Ionic Concentration Calculation, Native contact analysis, Root mean square fluctuation analysis, Fluctuation cross-correlation analysis, Principal Component Analysis, Residence Time Analysis; Supporting Results: Molecular interactions guiding the binding pathway, Fluctuation analysis of Tat peptide in the binding pathway, Residence Time Analysis for Sodium ions; Supporting Figures: molecular structure corresponding to each binding step, slope change in the free energy landscape, molecular interactions corresponding to each binding step, consistent dissociation pathway of TAR-Tat complex, backtracking event, dynamic cross correlation maps for apo TAR RNA and other binding step, PCA structural mode corresponding to bulge-tetraloop anticorrelation, time evolution of native contact of tetraloop and total contact for tetraloop fluctuation attenuated simulation, radial distribution function and residence time distribution of water around the TAR RNA, molecular picture of water mediated interaction at step B, radial distribution function of Tat peptide around TAR RNA and sodium around TAR RNA, residence time distribution of sodium ions around TAR RNA, overlap of each umbrella window along reaction coordinate, RMSD for equilibrium simulation of BIV TAR RNA and BIV TAR-Tat complex; Supporting Tables of slope associated with each binding step, no. of solvent and co-solvent added for system preparation, bulk ionic concentration of ions in the system, timescale of residence of sodium ion in step B, step A and apo form of TAR RNA; References. (PDF)

## AUTHOR INFORMATION

### AUTHOR CONTRIBUTION

Conceptualization: SR. Data curation: SP, RS. Formal analysis: SR, SP. Funding acquisition: SR. Methodology: SP, RS. Project administration: SR. Resources: SR. Validation: SR, SP. Writing – original draft: SR, SP. Writing – review & editing: SR, SP.

#### Notes

Authors declare no competing interests.

## ACKNOWLEDGEMENTS

SP, RS, and SR thank the DIRAC supercomputing facility at IISER-Kolkata for computational support. SR acknowledges support from the Department of Biotechnology (DBT) (Grant No. BT/12/IYBA/2019/12) and Science and Engineering Research Board (SERB), Department of Science and Technology (DST), Govt. of India (Grant No. SRG/2020/001295). SP acknowledges support from DST-INSPIRE scholarship for higher education(SHE).

## Data and materials availability

All data and codes used in the analysis are available from the corresponding author to any researcher for purposes of reproducing or extending the analysis under a material transfer agreement with IISER-Kolkata, India.

## TABLE OF CONTENTS GRAPHICS

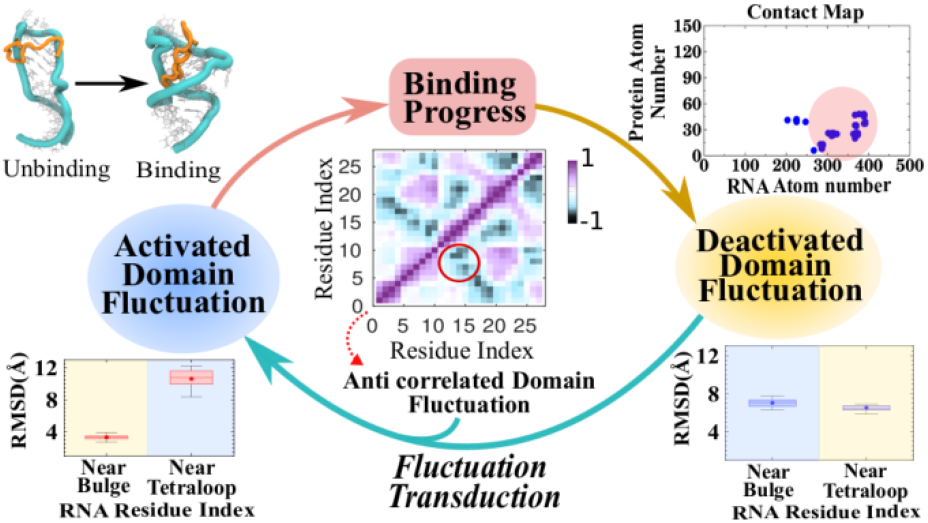

