## Supporting Information File for "Correlation in Domain Fluctuations Navigates Target Search of a Viral Peptide along RNA"


#### This PDF file includes:

**Supporting Information:**


- Supporting Methods:
  - Tetraloop Restrained Simulations
  - Bulk Ionic Concentration Calculation
  - Native contact analysis
  - Root mean square fluctuation analysis
  - Fluctuation cross-correlation analysis
  - Principal Component Analysis
  - Residence Time Analysis
- Supporting Results:
  - Molecular interactions guiding the binding pathway
  - Fluctuation analysis of Tat peptide in the binding pathway
  - Residence Time Analysis for sodium ions
- Supporting Figures:
  - Figure S1 to S21
- Supporting Table:
  - Table S1 to S4
- References for Supporting Information

### Supporting Methods

#### *Tetraloop Restrained Simulation*

In order to investigate the causality relationship between the fluctuation and binding the tetraloop fluctuation was attenuated artificially. For attenuation of tetraloop fluctuation, a position restraint with a force constant of 5000 kJ/mol/nm$^2$ was applied on the residue 15-20 of the TAR RNA. Subsequently, unbiased molecular dynaimcs simulation were done using a leapfrog integrator with a time step of 2 fs and an average temperature of 300 K were maintained using a Nose-Hoover thermostat[1,2] with a relaxation time of 0.5 ps. The initial structure for the simulation was extracted from the SMD trajectory when the RNA-protein complex was in state D conformation.

#### *Ionic Concentration Calculations*

Time-averaged bulk ionic concentrations of different ions are calculated by taking the time-averaged ratio between the number of bulk ions to that of the number of bulk water molecules multiplied by the molarity of pure water. All species which are present beyond 2 nm of the RNA are considered to be bulk species according to the radial distribution function as depicted in Figure S15.

$$[Na^+]^* = 55.51M\left\langle \frac{B_{Na^+}}{B_{H_2O}} \right\rangle \qquad [Cl^-]^* = 55.51M\left\langle \frac{B_{Cl^-}}{B_{H_2O}} \right\rangle$$

To this calculated raw concentrations, we applied an electroneutrality correction term to account for the residual variation from charge neutrality in the bulk of the solution, assuming a linear response of ion density with concentration due to weak electrostatic potential in the bulk regime[3].

$$[j] = [j]^*\left(1 - q_j \frac{\sum_i q_i [i]^*}{\sum_i q_i^2 [i]^*}\right)$$

$$[Na^+] = [Na^+]^*\left(1 - \frac{[Na^+]^* - [Cl^-]^*}{[Na^+]^* + [Cl^-]^*}\right) \qquad [Cl^-] = [Cl^-]^*\left(1 - \frac{[Na^+]^* - [Cl^-]^*}{[Na^+]^* + [Cl^-]^*}\right)$$

where $q_j$ is the charge of the ionic species, $[j]^*$ and $[j]$ denotes the raw concentration and the corrected concentration, respectively. The bulk ionic concentration of NaCl for both the system was found to be around 100mM, which is in the physiological ionic range. Details about the value of the ionic concentrations are given in Table S3.

#### ***Native Contact Analysis***

For the calculation of native contacts, we took a distance cutoff of 4.3Å, which corresponds to the second minima in the radial distribution function of heavy atoms of protein with respect to the heavy atoms of RNA(Figure S18). The native contacts were filtered out from the native structure of the complex that is from the 1BIV PDB structure. Subsequently, these native contacts were used as the basis for calculating the evolution of the total number of native contact as the binding progressed. Subsequently, the number of native contacts formed were normalized with the time-averaged number of native contacts present in the equilibrium simulation of the BIV TAR-Tat complex to obtain the fraction of native contacts formed. This native contact fraction analysis was done on several of the association trajectories. After that, the ensemble of all these native contact fraction analyses was plotted as a function of the distance of the center of mass of TAR RNA and the Tat peptide to study the pathway of formation of the BIV TAR-Tat complex. Subsequently, clustering was done using Ward-linkage algorithm[4] as implemented in the scikit-learn python library taking a total cluster number to be equal to four[5].

Thereafter, the contact map was also plotted to analyze the molecular interaction-based picture of the evolution of native contacts. The native contact map was calculated for different binding states by calculating the frequency of occurrence of that native contact on the trajectory corresponding to that particular binding state using the same cutoff distance of 4.3Å. Subsequently, only those native contacts were plotted in the contact map, which had a frequency of greater than 0.6.

##### ***Root Mean Square Fluctuation(RMSF) Analysis***

Root mean square fluctuation describes the time-averaged deviation of the position of atoms in a molecule with respect to the average structure.

$$RMSF_i = \sqrt{\frac{1}{\tau}\sum_{t_j=1}^{\tau}\left|r_i(t_j) - \langle r_i\rangle\right|^2}$$

where, $RMSF_i$ represents the RMSF of the *'i'*th particle, *'τ'* is the total time over which we have to calculate the RMSF and $r_i(t_j)$ represents the instantaneous position of the particle *'i'* at a time *'$t_j$'*, and $\langle r_i\rangle$ represents the average position of *'i'*th particle. The RMSF of TAR RNA is calculated taking into account only the phosphate atom of the RNA for three steps B, C , D and also for the native state A in the free energy profile using the 'gmx rmsf' module of GROMACS 2018.3[6] to compare the change in the fluctuation pattern of different nucleotides in the TAR RNA as the binding progress.

#### *Fluctuation cross-correlation analysis*

The correlation between the atomic fluctuations was calculated considering the phosphate(P) atoms as the representative of each nucleotide of the TAR RNA. These P-atoms are considered as a junction of an elastic network. These junctions fluctuate under the influence of their neighboring junctions. These fluctuations follow a Gaussian distribution[7]. Therefore,

$$f(\Delta \boldsymbol{r_{ab}}) = (N^*/\pi)^{3/2}.e^{(-N^*\Delta \boldsymbol{r_{ab}}^2)}$$

Here, $\Delta r_{ab}$ is the fluctuation in the distance between P-atom 'a' and 'b' and $N^*$ is the normalization constant. Thus, potential associated with the fluctuations of the P-atoms can be expressed as $V = \{\Delta \boldsymbol{r}^{\boldsymbol{T}}\}\Gamma\{\boldsymbol{\Delta r}\}$, where Δr represents an N-dimensional column vector formed by the fluctuations of the P-atoms,{ $\mathbf{\Delta r_1, \Delta r_2, \Delta r_3, \ldots\ldots\ldots\ldots \Delta r_N}$} and Γ represents Kirchhoff matrix.

$$\Gamma_{ab} = \begin{cases} -N^*, if\ a \neq b\ and\ r_{ab} \leq r_c \\ 0, if\ a \neq b\ and\ r_{ab} > r_c \\ -\sum_{a\neq b} \Gamma_{ab}, if\ a = b \end{cases}$$

where, $r_c$ is the non-bonded contact cutoff distance. So, the fluctuation correlation between two P-atoms 'c' and 'd' can be written as,

$$\langle \Delta \boldsymbol{r_c}.\Delta \boldsymbol{r_d}\rangle = \left(\frac{1}{Z}\right).\int \Delta \boldsymbol{r_c}.\Delta \boldsymbol{r_d}.e^{-V/kT}d\{\Delta \boldsymbol{r}\} = [\Gamma^{-1}]_{cd}$$

where, Z is the partition function. So, the cross-correlation between the atomic fluctuation of P-atoms can be found from the off-diagonal elements of the inverse of the Kirchoff's matrix[7]. These cross-correlations are then normalized with respect to the self correlations as :

$$C_{cd} = \frac{\langle \Delta \boldsymbol{r}_c . \Delta \boldsymbol{r}_d \rangle}{\sqrt{\left(\langle \Delta \boldsymbol{r}_c{}^2 \rangle \langle \Delta \boldsymbol{r}_d{}^2 \rangle\right)}}$$

The normalised cross-correlation values lie in the range of -1 to 1. The positive values of correlation indicates movement in the same direction that is positively correlated and the negative values represent anti-correlated motions.

#### *Principal Component Analysis*

Based on the cross-correlation obtained, the co-variance matrix was diagonalised with an orthonormal transformation matrix to obtain the eigen value and the corresponding eigen vectors for each of the binding steps. These eigen vectors are called the principle components which corresponds to the direction of internal motion of the RNA. Subsequently the vector component per atom(in this case the phosphate atoms only) of the eigen vectors were calculated to capture the magnitude and direction of motion of each P atom of the RNA with respect to each other. The principal component analysis for each of the binding step was performed using GROMACS 2018.3 to capture the evolution of correlation between the different domains of the RNA[8].

#### *Residence Time Analysis*

The residence time can be defined as the maximum amount of time a molecule/atom spends inside a particular cutoff around a reference point. In our case we calculated the residence time of water and sodium to explore their role in the binding of the TAR RNA with the Tat peptide. For calculating the residence time for water and sodium, a cutoff of 4.5Å and 5.0Å is taken, respectively. The cutoff is chosen based on the radial distribution function of water and sodium around the RNA as shown in Figure S15 and S19. For each water molecule and sodium ion, we calculated its distance from each of the atoms of RNA and a boolean variable is given to 1 if it satisfies the condition that the the species lies within its defined cutoff. But, if the species doesn't obey the condition, then a value of 0 is assigned to it[3,9]. This process is extended to all the water molecules and sodium ion present in the system for several time frames. Subsequently for each water molecule its maximum continuous stay within the cutoff is taken as its residence time. This residence time is calculated for free TAR RNA, native state of TAR-Tat complex(A) and state B during Tat peptide binding to TAR RNA.

#### Supporting Results

##### *Molecular Interactions stabilizing and guiding the binding pathway*

The very first interaction for the binding of BIV-TAR RNA with Tat peptide is initiated by Arg32 and Arg34, as shown in Figure S4. Due to the presence of the guanidium group, arginine can form hydrogen bonds with the phosphate group of the RNA. These bonds are electrostatically also stabilized as the guanidium group contains a positive charge and the phosphate groups contain a negative charge on the oxygen atoms. Thus this interaction between the guanidium group of Arg32 and Arg34 with the oxygen of nitrogenous base of C14 and phosphate group of A18 respectively forms the first interaction site for TAR-Tat complex formation.

This initial binding of Arg32 and Arg34 by hydrogen bonding leads to the next stable interaction in the binding pathway where Arg32 and Arg34 are involved in more stable interactions, namely base stacking, and base triple, as shown in Figure S5. These interactions are the signature of state D. In this interaction, Arg32 begins to form the first base triple interaction between Arg32, A18, and G19. This interaction provides the binding pathway its first most stable anchoring point as this base triple is present in the native structure. Moreover, Arg34 also interacts with G11 and C20 to form a strong base triple pairing interaction, which also propagates till the native structure. So, this two interactions are the first native contacts formed in the binding pathway[10].

In state D, the base stacking between Arg32, A18, and G19 begins to form, and in state C this Arg32, A18, and G19 base stacking reach its stable conformation. This conformation is present in the native TAR-Tat complex. Moreover, the base triple interaction between Arg34, G11, and C20 is also preserved at this point in the binding pathway as shown in Figure S6. The new interaction which comes into play at this point is the hydrogen bonding interaction between Thr36 and C20. This interaction occurs at two points- between (i)carbonyl oxygen of Thr36 and the amino group of C20, (ii)hydrogen of the hydroxyl group of Thr36 and the phosphate link between C20 and U21. This interaction is also present in the native structure. Patel and co-workers[10] have also discovered this interaction in the native TAR-Tat complex structure.

In state B, some of the strong interactions between the RNA and the protein are preserved, such as the interaction between Arg32-A18-G19, C20-G11-Arg34, C20-Thr36. Besides this, a new non-native contact between Lys39, G6, and C23 also comes into play, as shown in Figure S7. This interaction is somewhat similar to that of arginine hydrogen bonding interaction as Lys is also a positively charged amino acid because of a protonated amine group in its side chain. Hence, the Lys39 interacts through its side chain amino group with G6 and C23 to form a triple base pairing interaction(Figure S7).

The formation of interaction present in state B helps the TAR-Tat complex to achieve its most stable structure that is the native structure. When the complex is in minima B most of its native contact has been formed in each step of binding preceding this last step, such as Arg32-A18-G19, C20-Thr36 (formed at minima C), and C20-G11-Arg34( formed at minima D)[10,11]. The last most crucial rearrangement which occurs during the transition from state B to the native state A is the change in interaction from Lys39-G6-C23 to Arg41-G6-C23. This change in interaction is energetically favorable due to the presence of the guanidium group in arginine, which has a more substantial hydrogen bonding interaction potential than that of lysine. Thus this interaction leads to a very stable base triple interaction between Arg41, G6, and C23, as shown in Figure S8.

So, from these observations about the molecular interactions governing the pathway of binding of the TAR RNA to the Tat peptide, we can say that Arginine is playing an essential role in this process; hence the arginine-rich motif is crucial for this binding. Various other research groups have also made a similar observation[10–13].

#### *Fluctuation Analysis of the Tat peptide in the binding pathway*

In this study, the Tat peptide used in simulations is a small highly intrinsically disordered peptide (17 mer). To investigate the role of the Tat peptide in the binding pathway we have calculated the RMSF of the Tat peptide at each of the binding states. The RMSF has been calculated on the Cα atom of each amino acid in the peptide for clear analysis.

It was observed that as the binding initiates, the peptide shows higher fluctuation, and as the first binding happens, there is a complete supression in the fluctuation as shown in Figure S12. Moreover, due to its disordered nature, there is an absence of any particular trend in the fluctuation of each residue as the binding progress and it lacks the domain-domain relationship as present in the case of TAR RNA. Thus, the fluctuation in the TAR RNA becomes more essential in this binding process.

##### *Residence Time Analysis for Sodium ions*

It has been observed from the residence time plot (Figure S20) that for both state B and state A the mean residence time is nearly the same that is around 1ns. However, for free BIV TAR RNA the mean residence time of sodium ions are higher than that of state B and state A. Thus, the sodium ion is not playing a very dominant role in stabilising the native state interactions as evident from both Figure S20 and Table S4.

### Supporting Figures

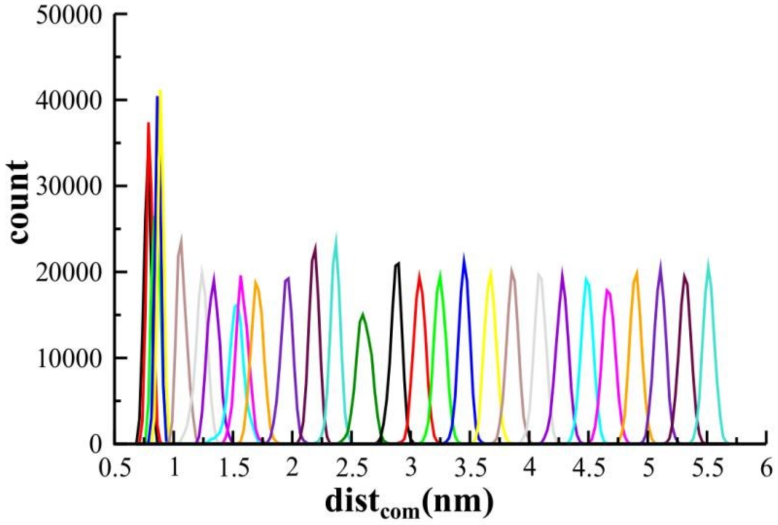


**Figure S1.** Overlap of the probability distribution of each umbrella windows spread through the reaction coordinate.

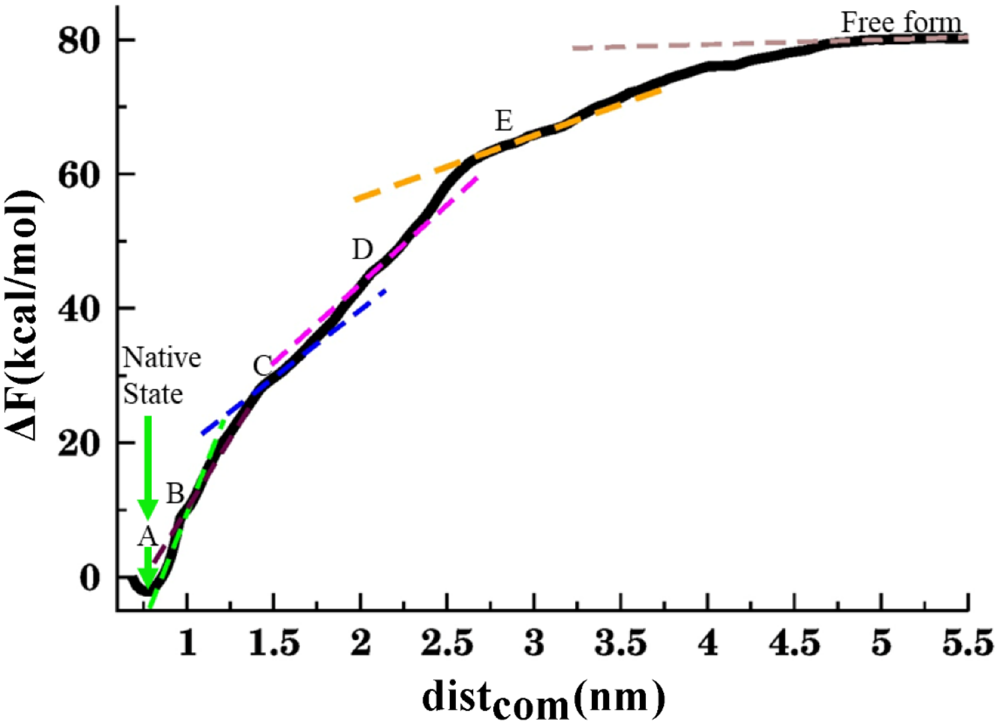


**Figure S2**. Free Energy profile with metastable humps characterised by change in slope of barrier.

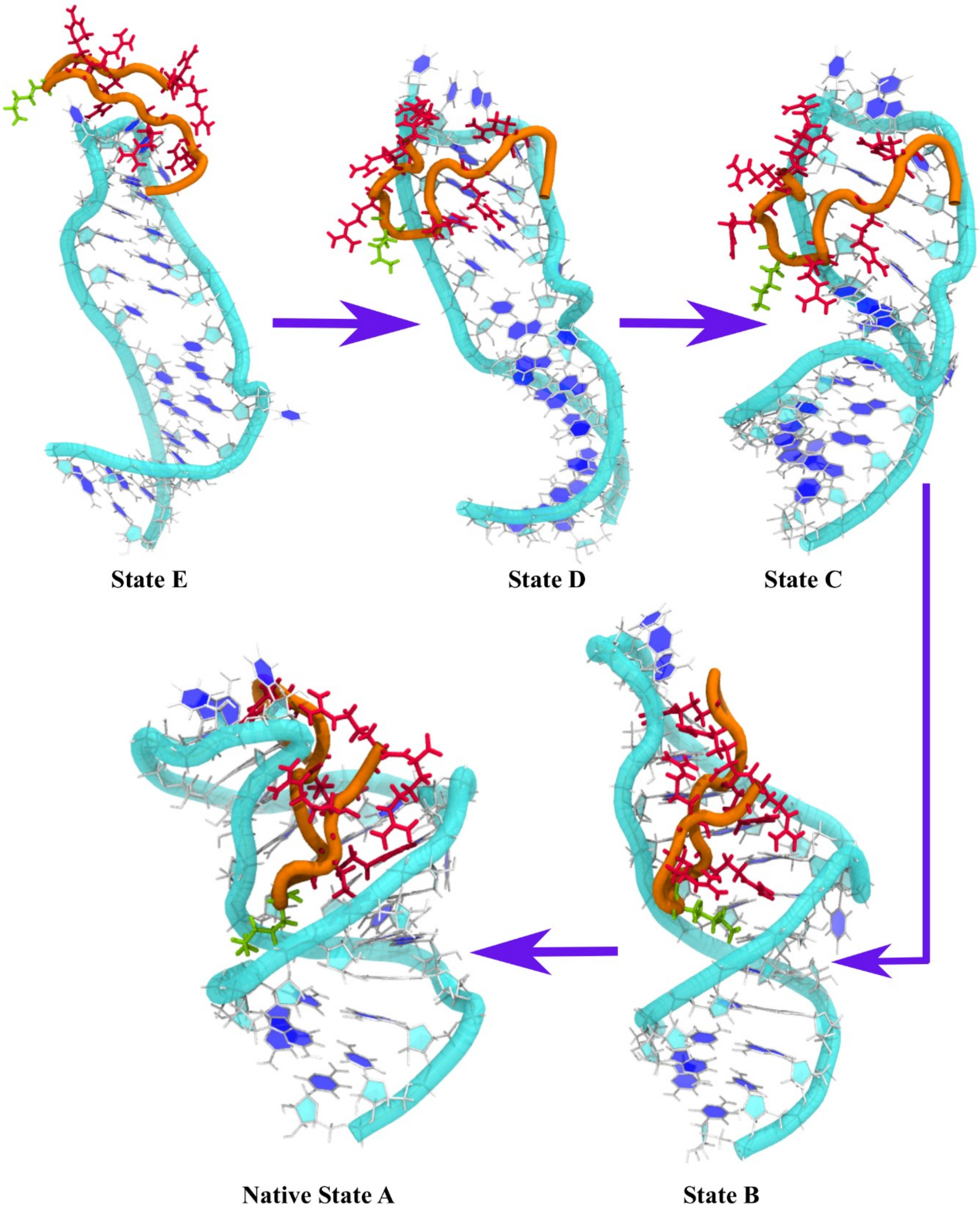


**Figure S3**. Molecular structure of BIV TAR-Tat conformation corresponding to the subsequent binding state associated with the TAR-Tat binding pathway.

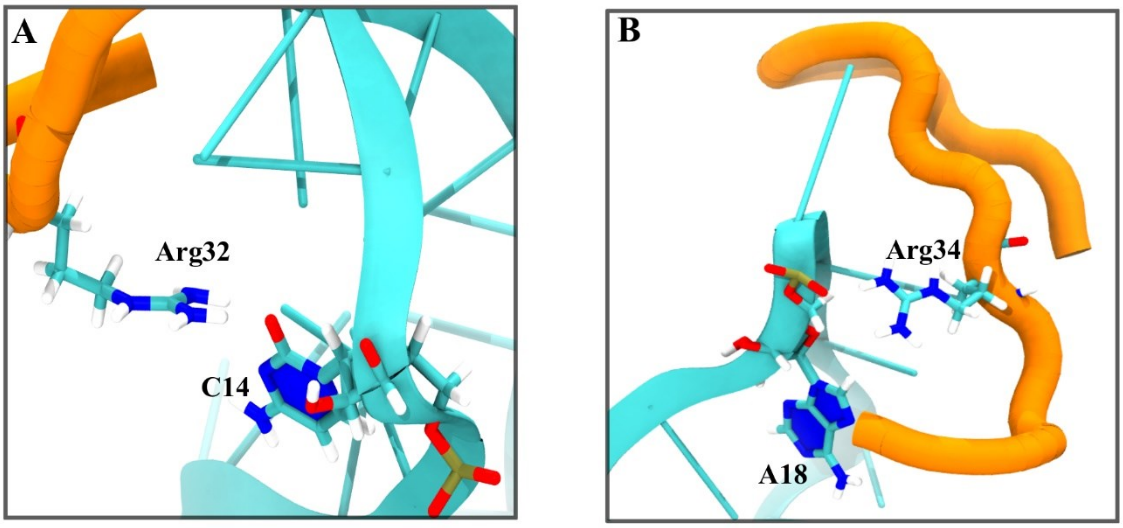


**Figure S4**. Interactions near the tetraloop initiating the first interaction between TAR RNA and Tat peptide(minima E). (A)interaction between Arg32 guanidium group and oxygen of nitrogenous base of C14. (B) Guanidium group of Arg34 interacting with phosphate oxygen of A18.

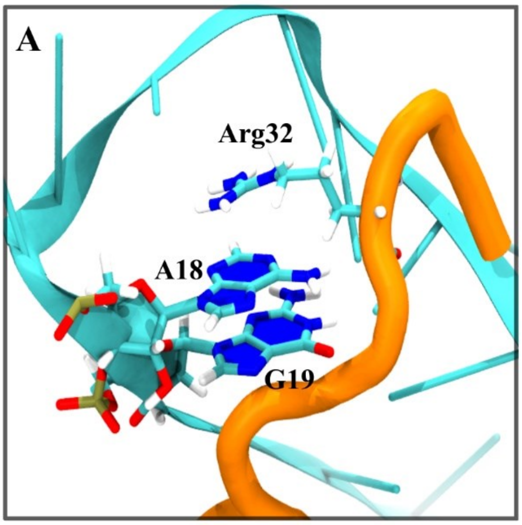


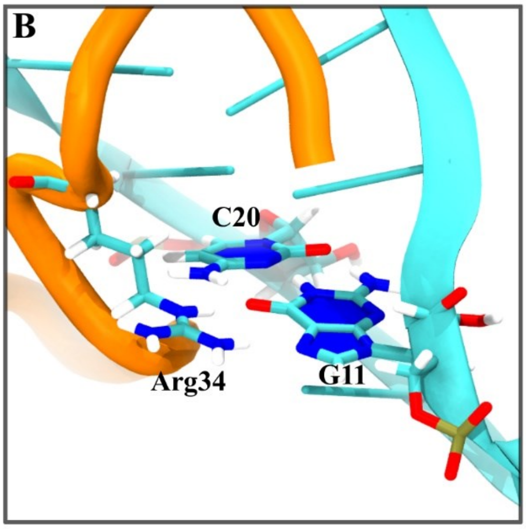


**Figure S5**. Interactions stabilizing minima D. (A)base stacking interaction between Arg32, A18, G19. (B) base triple interaction between Arg34, G11, C20

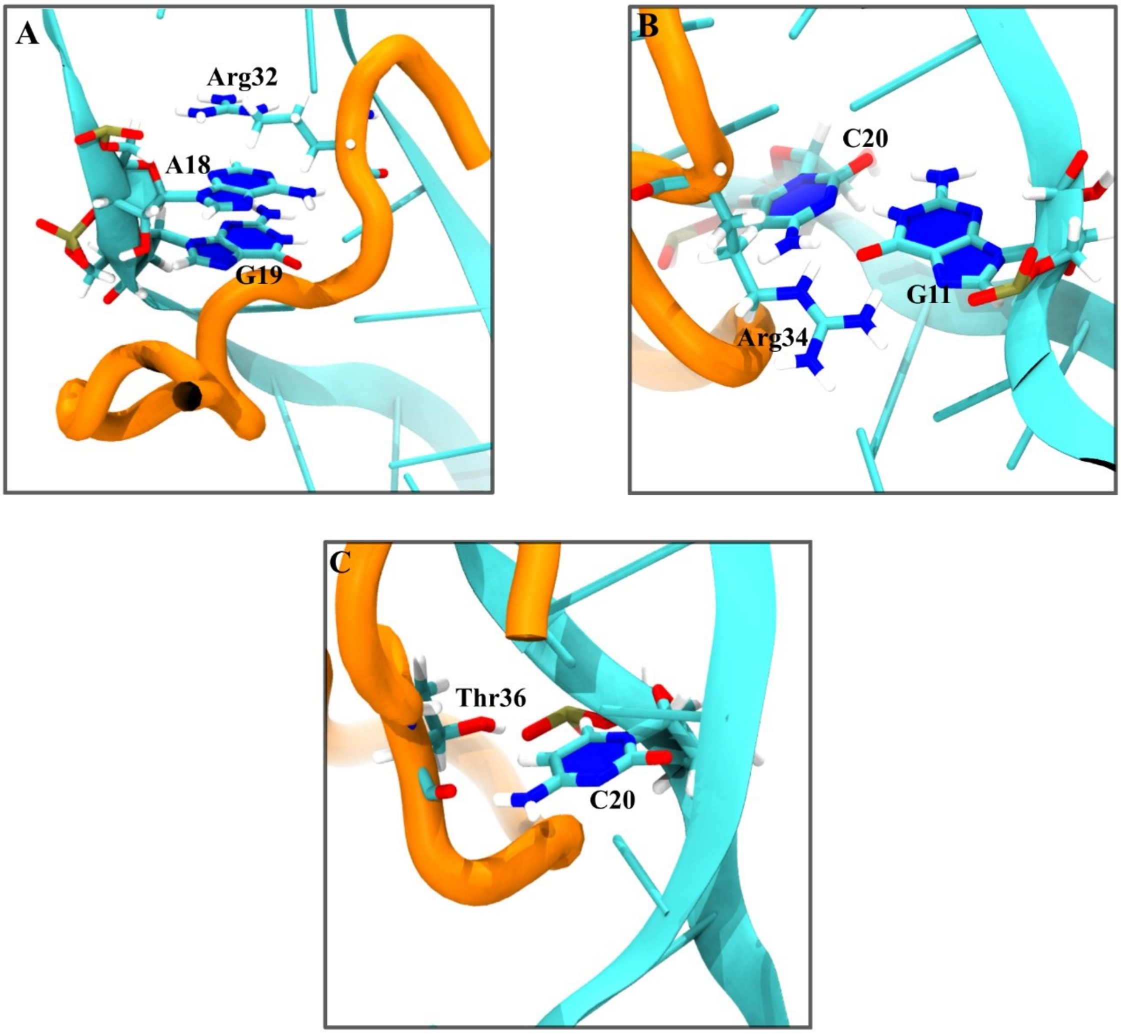


**Figure S6**. Interactions stabilizing state C. (A)base stacking interaction between Arg32, A18, G19 (B) base triple interaction between Arg34, G11, C20 (C) hydrogen bonding interaction between Thr36 and phosphate group joining C20-U21

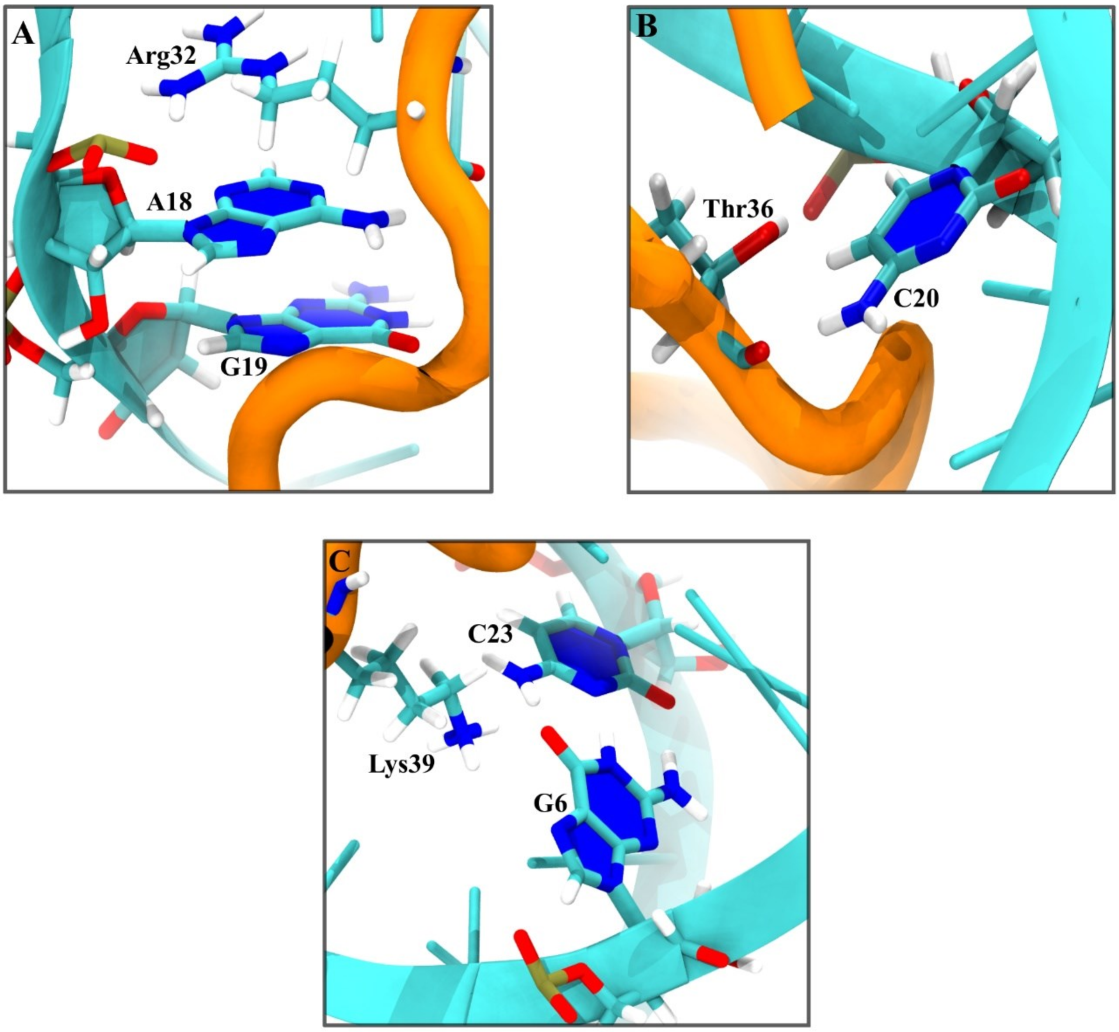


**Figure S7.** Interactions stabilizing state B. (A)base stacking interaction between Arg32, A18, G19 (B) hydrogen bonding interaction between Thr36 and phosphate group joining C20-U21 (C) base triple interaction between Lys39, G6 and, C23.

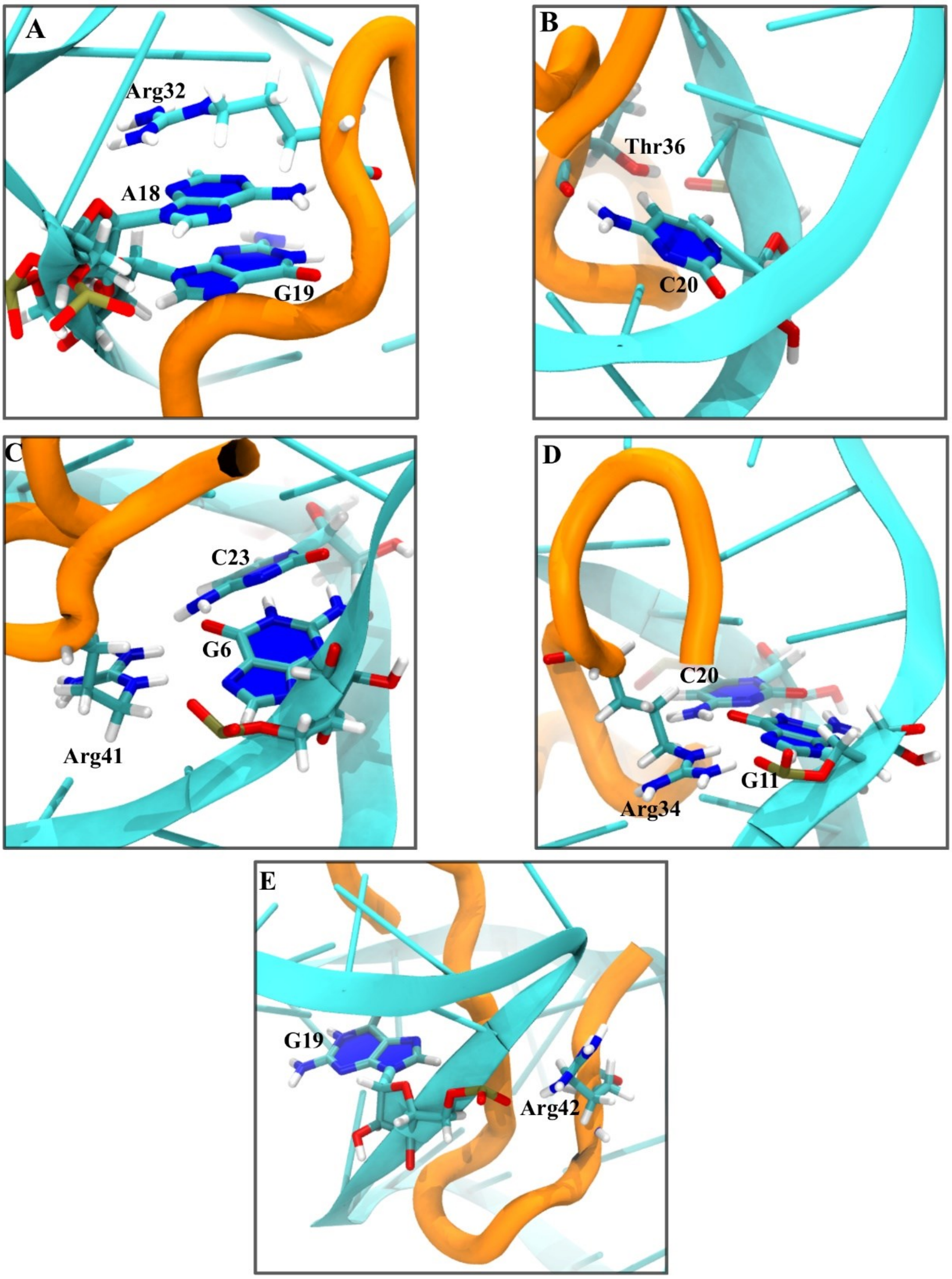


**Figure S8**. Interactions stabilizing the native structure of TAR-Tat complex. (A)base stacking interaction between Arg32, A18, G19 (B) hydrogen bonding interaction between Thr36 and phosphate group joining C20-U21 (C) base triple interaction between Arg41,G6 and C23 (D) base triple interaction between Arg34, G11, C20 (E) backbone interaction between the phosphate group of G19 and the Arg42.

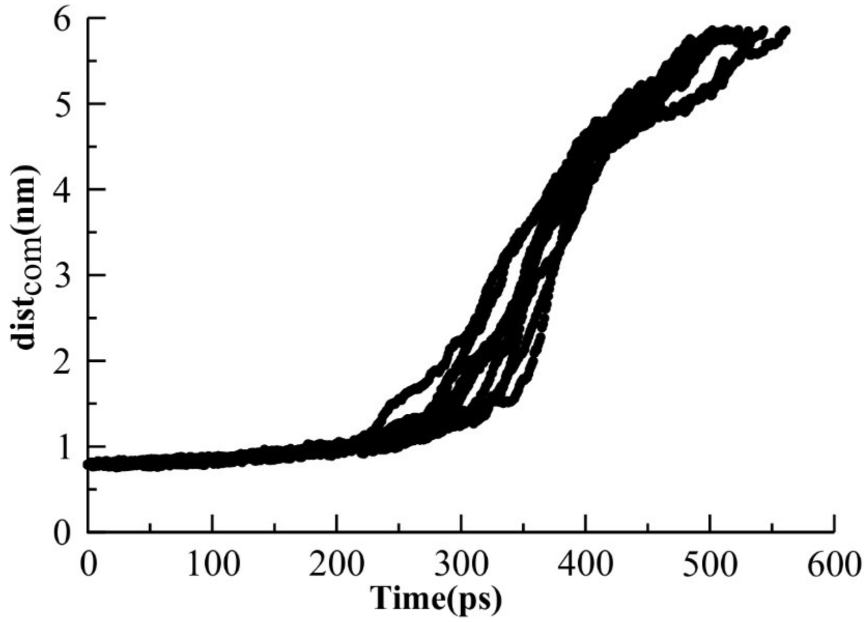


**Figure S9**. Consistent pathway of dissociation of Tat peptide from the TAR RNA in SMD simulations. Plot of $dist_{com}$ against time as the protein is pulled from the RNA in 10 different constant force pulling simulations.

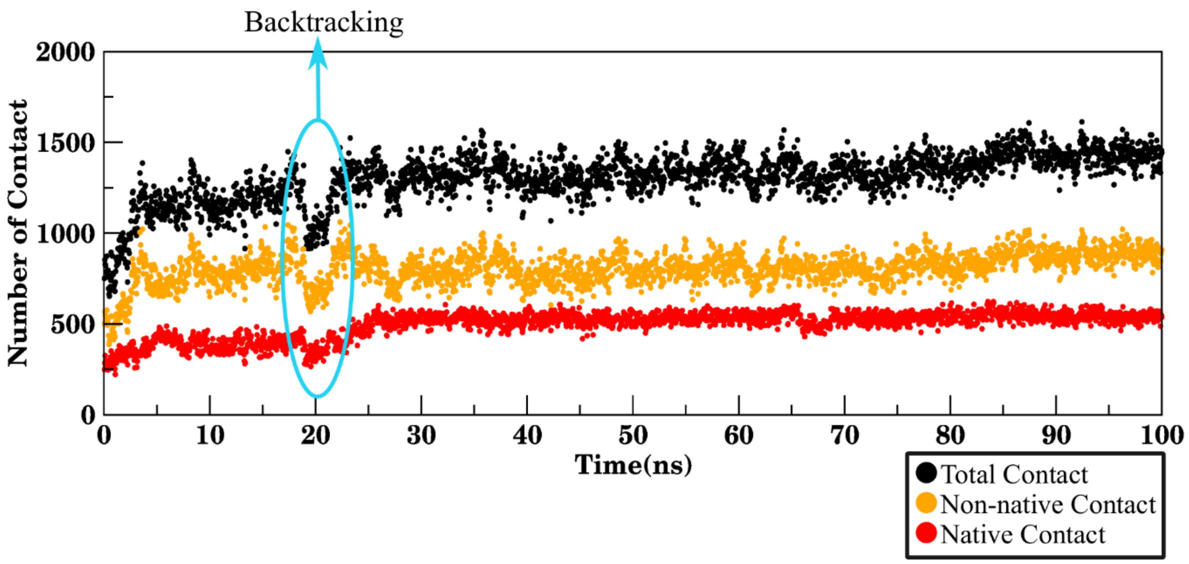


**Figure S10**. Evolution of number of native contact(red) , non-native contact(orange) and total contact(black) in a 100ns simulation. The blue circle shows the backtracking event captured in some of the simulations.

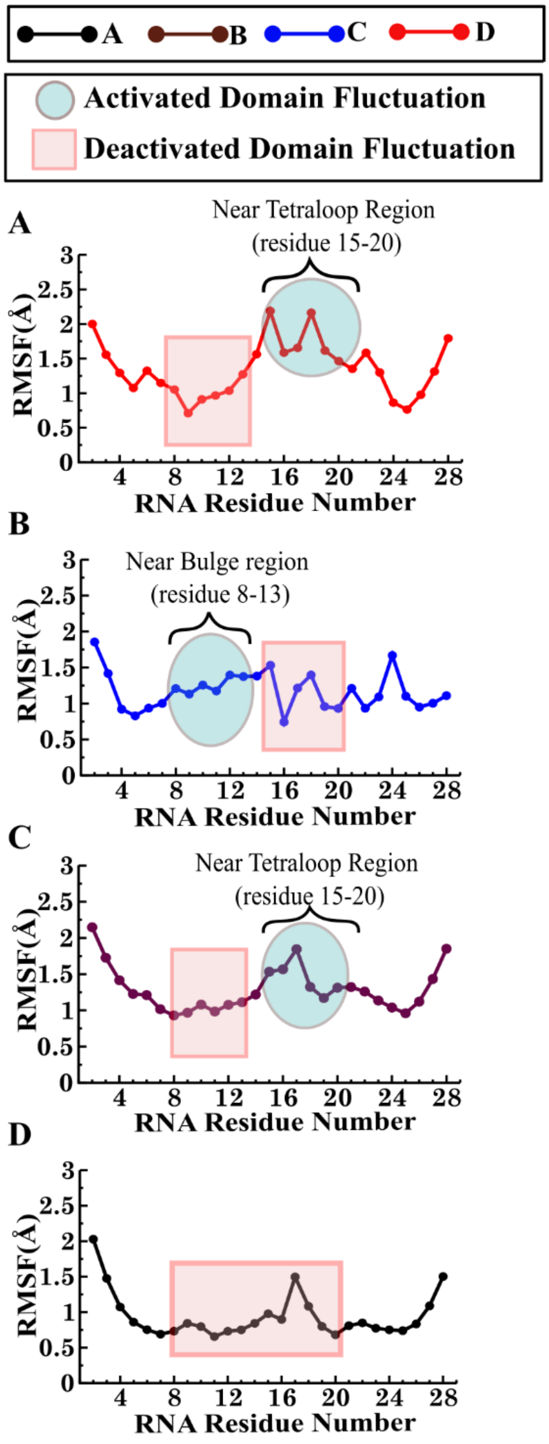


**Figure S11**. Root mean square fluctuation(RMSF) of TAR RNA of (A) state D (B) state C (C) state B (D) state A of binding pathway calculated on phosphate atom of each nucleotide.

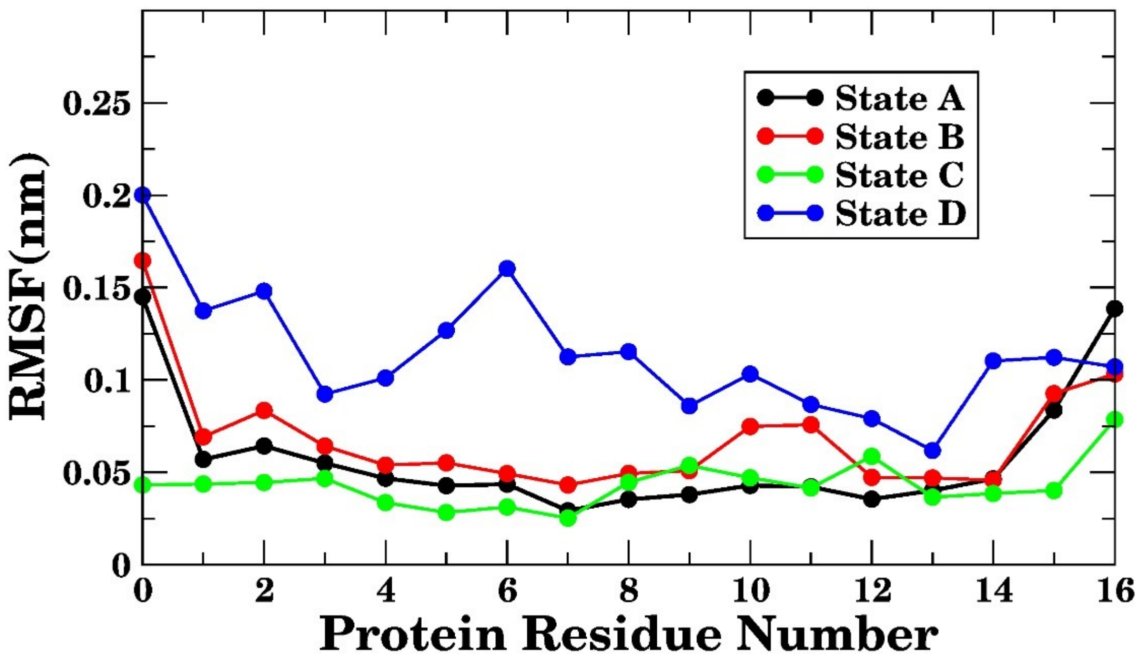


**Figure S12**. Root mean square fluctuation(RMSF) of Tat peptide for binding state D, C, B and A calculated on each Cα-atom of each amino acid of Tat peptide

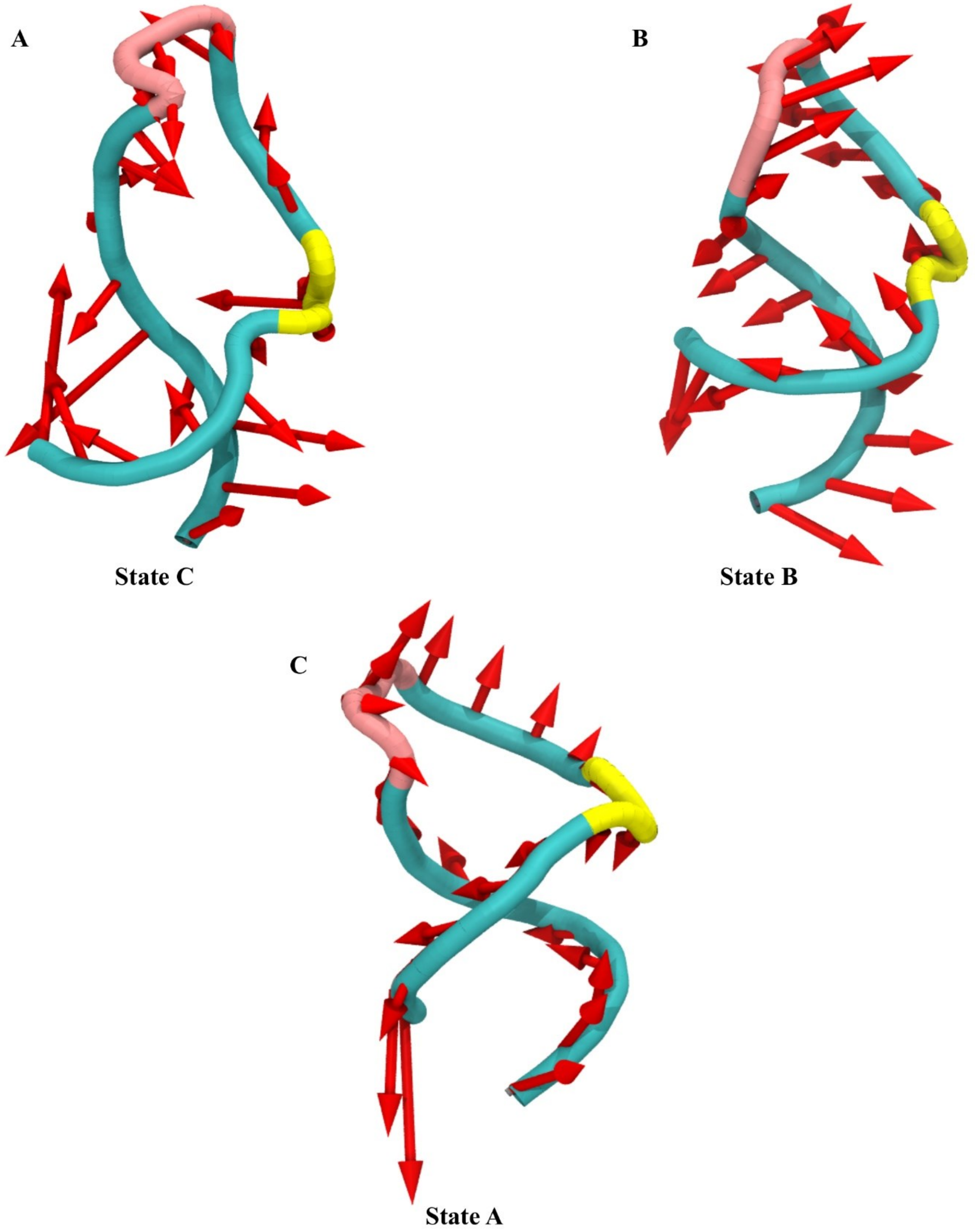


**Figure S13**. Slow bulge-tetraloop anti-correlated motion modes of TAR RNA corresponding to binding state C, B and A. The TAR RNA backbone is shown as a cyan colored tube with tetraloop and bulge shown in pink and yellow colour respectively. The modes are shown with red coloured arrows.

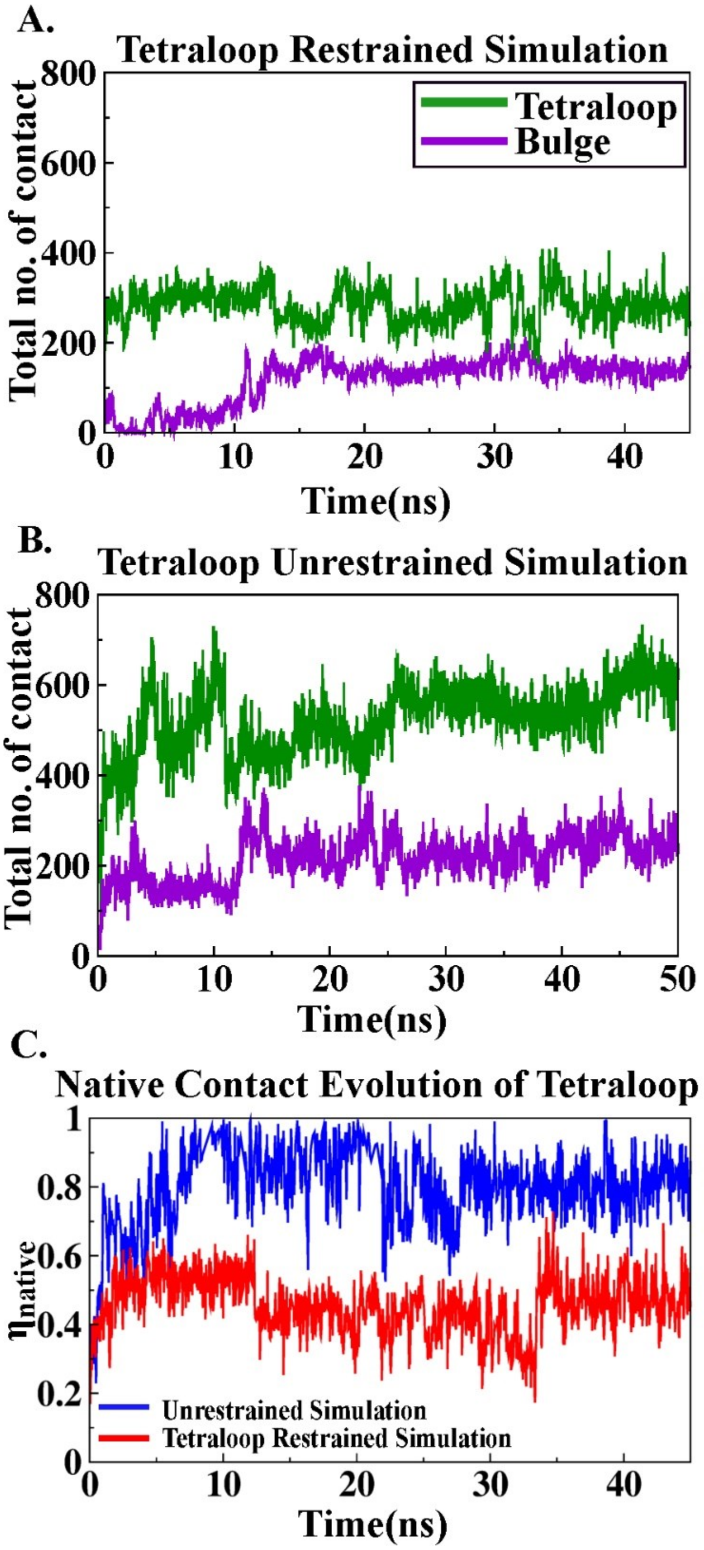


**Figure S14**. (A)Time Evolution of total contact for near tetraloop restrained unbiased simulation (B) Time Evolution of total contact for near tetraloop unrestrained unbiased simulations. (C) Time Evolution of native contact of tetraloop for unrestrained and restrained simulation.

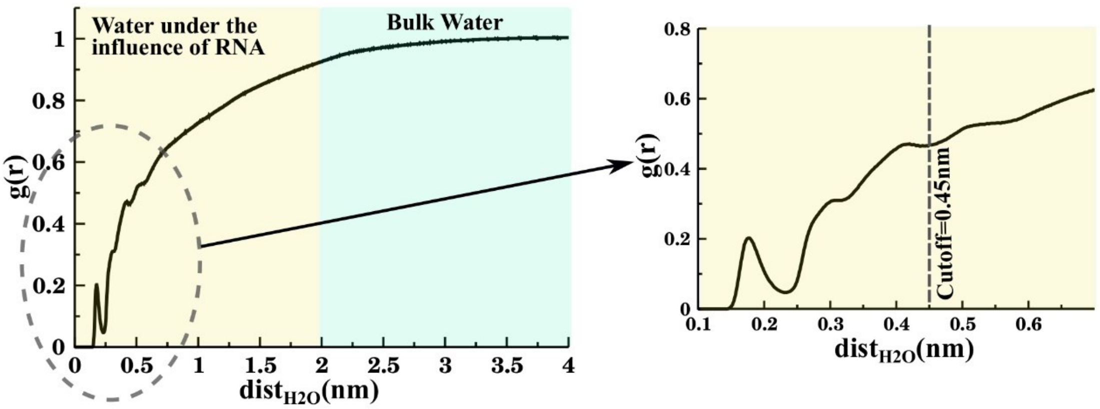


**Figure S15.** Radial Distribution function of water with respect to TAR RNA taking into account all non-hydrogen atoms in both TAR RNA and water. The yellow shaded region implies the distance upto which there is a large scale influence of TAR RNA on the water distribution and the light green shaded region implies the bulk water which either doesnt have a significant influence of the TAR RNA.The black dotted line in the inset figure shows the cutoff considered for calculation of residence time of water.

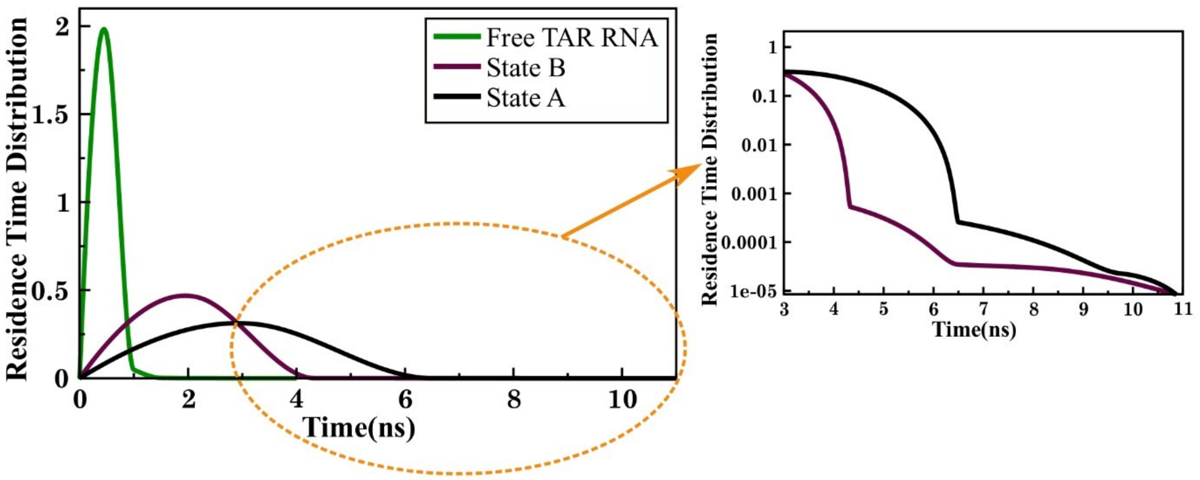


**Figure S16.** Residence time distribution of water molecules around the TAR RNA for free TAR RNA(green), state B of TAR-Tat complex(maroon) and native state of TAR-Tat complex(black). The residence time distribution of the water molecule which have a stay period of greater than 3 ns within the cutoff is highlighter in the inset figure with the residence time being logged.

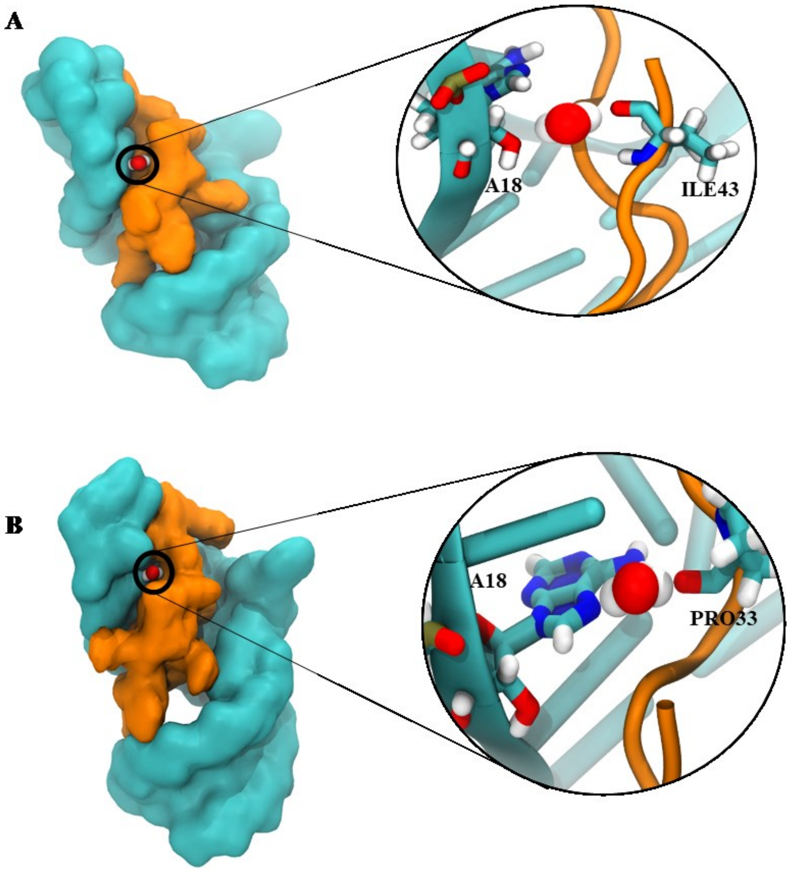


**Figure S17.** Highly residing water molecules bridging the TAR-Tat interaction in state B of the binding pathway. (A) One of the highly residing water molecules is mediating the interaction between the phosphate group of adenine and the carboxyl group of isoleucine. (B) The nitrogen of the nucleotide base of the adenine interacts with the proline carboxyl group with the help of a highly residing water molecule.

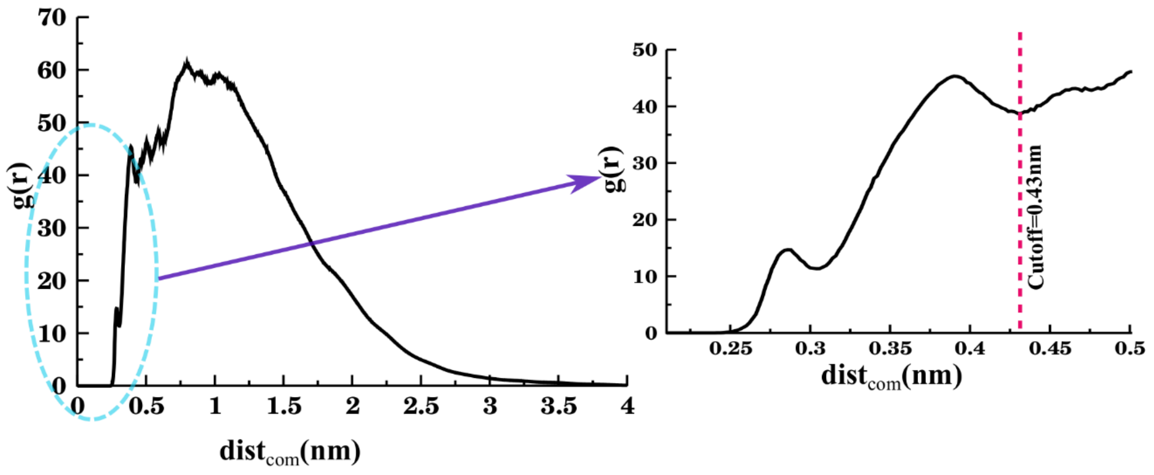


**Figure S18**. Radial Distribution function of Tat peptide with respect to TAR RNA taking into account all non-hydrogen atoms in both TAR RNA and Tat peptide. The red dotted line in the inset figure shows the cutoff considered for calculation of native contacts

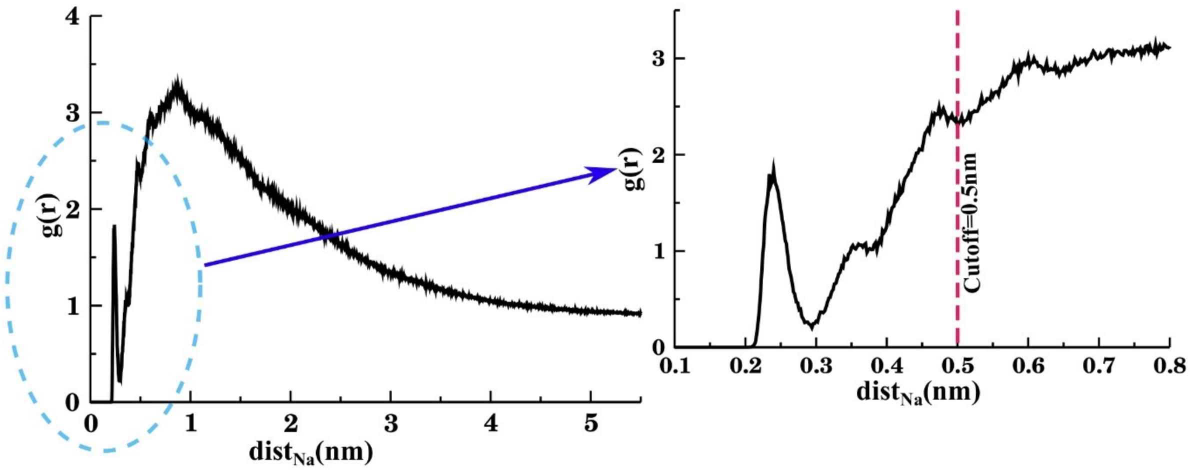


**Figure S19**. Radial Distribution function of sodium(Na) ion with respect to TAR RNA considering all non-hydrogen atoms in TAR RNA. The red dotted line in the inset figure shows the cutoff considered for calculation of residence time of Na ions.

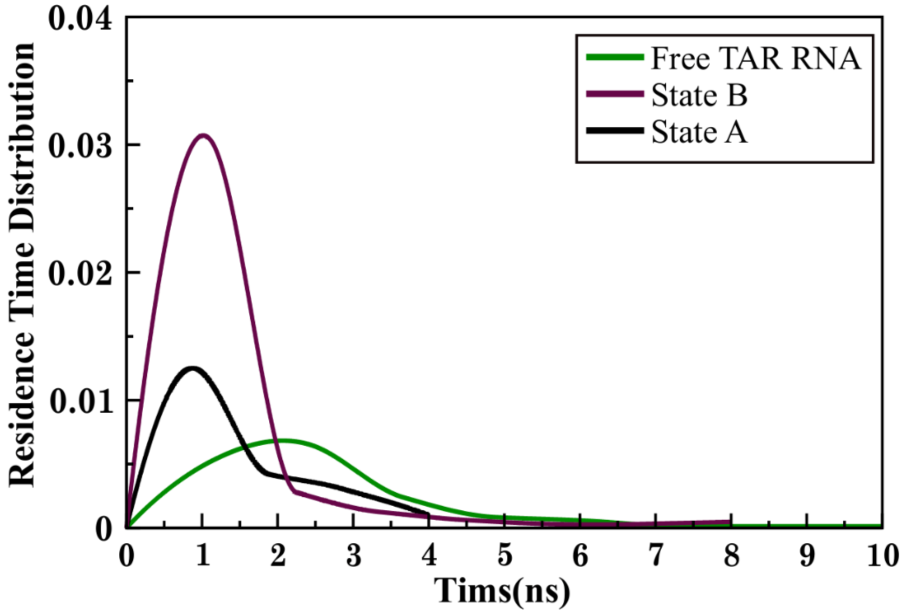


**Figure S20.** Residence time distribution of sodium ions around the TAR RNA for free TAR RNA(green), state B of TAR-Tat complex(maroon) and native state of TAR-Tat complex(black). It is observed that the mean residence time of sodium ions in free TAR RNA is greater than that of the TAR-Tat complexed form.

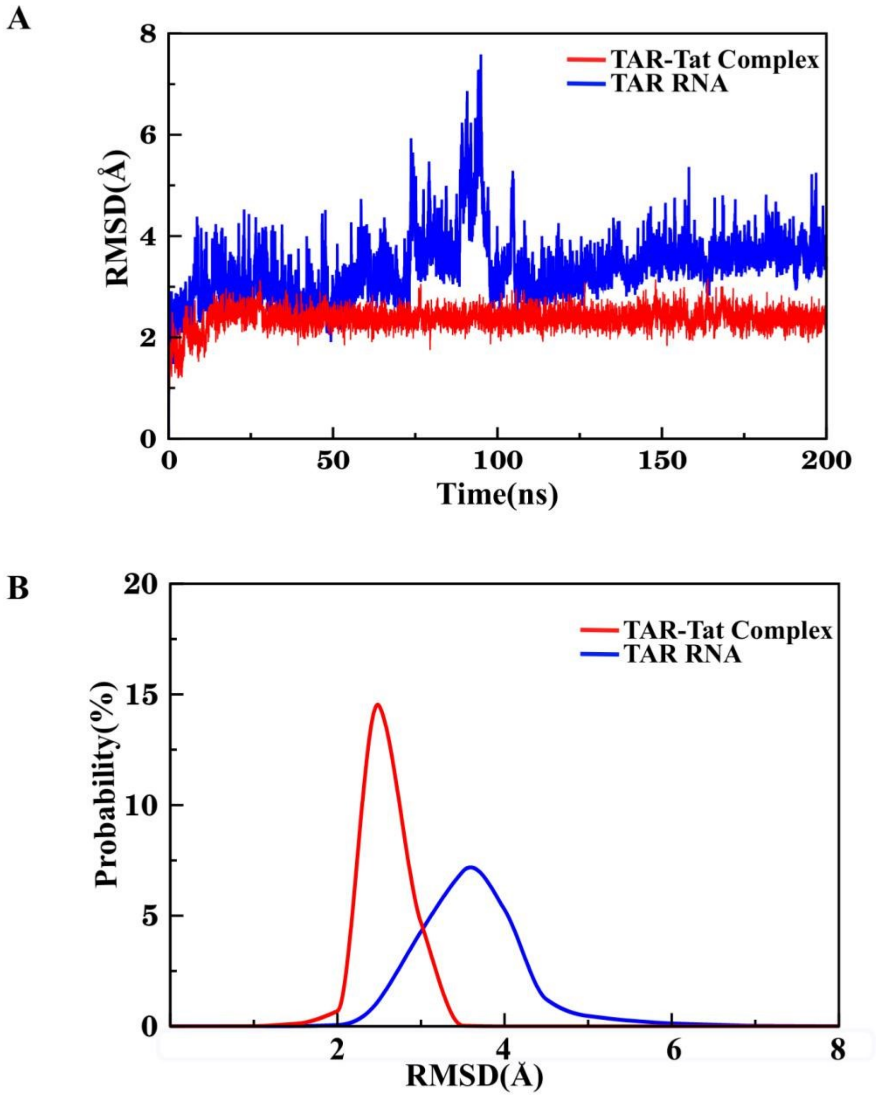


**Figure S21.** (A) Root mean square deviation of equillibrium MD simulation trajectory for BIV TAR-Tat complex and free BIV TAR RNA. (B)Probability distribution of RMSD calculated on the unbiased molecular dynamics simulation trajectories.

#### Supporting Tables

**Table S1:** Number of solvent and co-solvent added to BIV TAR RNA system and BIV TAR-Tat complex system to maintain the charge neutrality and to mimic the physiological ionic environment of the RNA and RNA-protein system.

| | No. of water molecule | No. of Sodium ion | Number of Chloride ion |
|---|---|---|---|
| BIV TAR RNA | 56308 | 116 | 89 |
| BIV TAR-Tat Complex | 55974 | 116 | 97 |

**Table S2:** Slopes associated with different steps of binding in the free energy landscape.

| **Steps** | **Slope of fitted straight line** |
|---|---|
| Completely Dissociated Protein | 0.72 |
| First binding of protein(E) | 9.28 |
| Second Step of binding(D) | 23.52 |
| Third Step of binding(C) | 20.21 |
| Pre-Final binding step Minima B | 42.03 |
| Final binding step to reach Native State A | 65.52 |

**Table S3:** Calculation of bulk concentration of ions in both TAR RNA system and TAR-Tat complex system.

| | Ions | No. of bulk ion | No. of bulk water | Rough Concentration | Corrected Concentration |
|---|---|---|---|---|---|
| BIV TAR RNA | $Na^+$ | 91.36 | 49962.45 | 101.5mM | 99.4mM |
| | $Cl^-$ | 87.67 | 49962.45 | 97.4mM | 99.4mM |
| BIV TAR-Tat Complex | $Na^+$ | 93.45 | 49781.53 | 104.3mM | 101.9mM |
| | $Cl^-$ | 89.46 | 49781.53 | 99.75mM | 101.9mM |

**Table S4:** The table categorizes the sodium ions into three categories: sodium ions staying for (i) >6ns (ii) 4-6ns (iii) 2-4ns. Based on these three categories, the number of sodium ions found in each category at three different states of the BIV TAR-Tat system(free TAR RNA, Step B, and native state A) is recorded in the table. It was found that as we move from free BIV TAR RNA to step B and subsequently to native state A, the number of sodium ions having high residing time decreases.

| | Number of Na ions staying for | | |
|---|---|---|---|
| System | >6ns | 4-6ns | 2-4ns |
| State A TAR-Tat Complex | 0 | 0 | 20 |
| State B TAR-Tat Complex | 1 | 1 | 12 |
| Free TAR RNA | 4 | 9 | 34 |
